# Androgen Signaling Restricts Glutaminolysis to Drive Sex-Specific Th17 Metabolism

**DOI:** 10.1101/2023.02.10.527741

**Authors:** Nowrin U Chowdhury, Jacqueline-Yvonne Cephus, Matthew Z Madden, Melissa M Wolf, Channing Chi, Ayaka Sugiura, Matthew T Stier, Kelsey Voss, Xiang Ye, Shelby N Kuehnle, Kennedi Scales, Vivek D Gandhi, Robert D. Guzy, Katherine N Cahill, Anne I Sperling, R. Stokes Peebles, Jeffrey C Rathmell, Dawn C Newcomb

## Abstract

Females have increased prevalence of many Th17-mediated diseases. While androgen signaling decreases Th17-mediated inflammation, the mechanisms are not fully understood. Th17 cells rely on glutaminolysis; however, it remains unclear whether androgen receptor (AR) signaling in males modifies glutamine metabolism to suppress Th17-mediated inflammation. We show that Th17 cells from male humans and mice had decreased glutaminolysis compared to females, and AR signaling attenuated Th17 cell mitochondrial respiration and glutaminolysis.

Using allergen-induced airway inflammation models, we determined females, but not males, had a critical reliance upon glutaminolysis for Th17-mediated airway inflammation, and AR signaling attenuated glutamine uptake by reducing expression of glutamine transporters. These findings were confirmed in circulating human Th17 cells with minimal reliance on glutamine uptake in male compared to female Th17 cells. We found that AR signaling attenuates glutaminolysis, demonstrating sex-specific metabolic regulation of Th17 cells with implications for design and implementation of Th17 or glutaminolysis targeted therapeutics.

**Highlights:**

- Human male CD4+ T cells have decreased expression of metabolic enzymes and decreased reliance on glutaminolysis compared to female CD4+ T cells.
- Androgen signaling decreased mitochondrial metabolism in Th17 cells and decreased airway inflammation.
- Androgen signaling decreased glutamine uptake and utilization in Th17 cells.

## INTRODUCTION

Cellular metabolism drives CD4+ T cell subset differentiation and stability, and CD4+ T cell metabolic pathways are promising potential therapeutic targets for cancers, autoimmune diseases, and asthma (Buck et al., 2015; Feng et al., 2022; Wei et al., 2017). Once activated, effector CD4+ T cell subsets, including Th1, Th2, and Th17 cells, substantially increase glycolysis to proliferate and allow for clonal expansion (Buck *et al*., 2015; Michalek et al., 2011). However, the reliance on other metabolic pathways varies between Th1, Th2, Th17, and regulatory T cells (Tregs) (Gerriets et al., 2015; Michalek *et al*., 2011; Puleston et al., 2021; Wei *et al*., 2017). Prior research has heavily focused on the metabolic pathways required for Th1, Th17, and Tregs - as these CD4+ T cell subsets are critical in pathogenesis of cancers and autoimmune diseases. Here, we focus on the metabolic pathways that drive Th17 and Th2 cells.

While an early increase in glycolysis is essential for activation of effector CD4+ T cell subsets, recent studies have focused on the differential metabolic pathways essential for subset-specific function. The metabolic dependencies of Th2 cells have been limited, but studies have found that Th2 cells also rely upon fatty acid oxidation and PPAR-γsignaling (Stark et al., 2019).

Exploration of Th17 metabolism has revealed several dependencies. Th17 cells require de novo fatty acid synthesis (Berod et al., 2014) and one carbon metabolism (Sugiura et al., 2022) for differentiation, effector function, and cytokine expression (Kono, 2022). Pathogenic Th17 cells have recently been shown to rely upon OXPHOS and mitochondrial metabolism (Baixauli et al., 2022; Buck et al., 2016; Hong et al., 2022; Shin et al., 2020). Further, glutaminolysis is essential for Th17 differentiation and effector function, important for Th2 function, and conversely may inhibit Th1 differentiation (Healey et al., 2021; Johnson et al., 2018; Kono, 2022; Kono et al., 2018). Glutaminolysis involves the uptake and conversion of glutamine to glutamate by glutaminase (GLS). Glutamate can then be converted to α-ketoglutarate which can enter the tricarboxylic acid cycle, be converted to glutathione for reactive oxygen species management, modify chromatin methylation patterns, or be used for other metabolic processes (Feng *et al*., 2022; Nakajima and Kunimoto, 2014; Siska et al., 2016; Wang et al., 2010)..

While metabolic needs for Th1, Th2, and Th17-mediated diseases have been previously studied, it remains unexplored how sex hormones modify these metabolic pathways.

Understanding the role of sex hormones is critical since there is a female predominance in the prevalence and/or severity of many immune-mediated diseases, including asthma, lupus, multiple sclerosis, rheumatoid arthritis, and type I diabetes (Klein and Flanagan, 2016). This female sex bias has been attributed to a myriad of factors, including impact of sex hormones, X and Y chromosomal differences, and social and environmental factors (Chowdhury et al., 2021; Klein and Flanagan, 2016), but the role of immunometabolism is unclear.

Prior research on sex hormone signaling and cell metabolism has shown in cancer that androgens signaling through the AR dysregulate glycolysis, the citric acid cycle, amino acid metabolism, and fatty acid metabolism through transcriptional, epigenetic, and co-receptor changes in prostate cancer cells (Barfeld et al., 2014; Massie et al., 2011; Uo et al., 2020). Further, AR signaling increased glutamine metabolism and utilization in gliomas (Sponagel et al., 2022). However, these studies have focused on cancer cells, which can have dysregulation of normal metabolic patterns and dysregulation of sex hormone receptor expression.

Additionally, various studies have described the impact of AR signaling on immune cells and T cells through reductions in function or increased expression of exhaustion markers but have not addressed the impact of sex hormones on immunometabolism (Fuseini et al., 2018; Gandhi et al., 2022; Kwon et al., 2022; Yang et al., 2022). Defining how sex hormones affect CD4+ T cell metabolism would provide a foundational mechanism for sex differences in CD4+ T cell subset effector function.

A previous study from our group showed that patients with asthma had increased numbers of CD4+ effector T cells and increased expression of metabolic enzymes in CD4+ T cell compared to healthy controls (Healey *et al*., 2021). However, the sex of participants and the impact of sex hormones on these cells was not considered. Given the importance of distinct metabolic programs on T cell function, we sought to determine the role of AR signaling in modifying CD4+ T helper cell metabolism *in vitro* using human and mouse cells and *in vivo* using a mouse model of allergic airway inflammation. Here, we showed that AR signaling directly decreased glutamine uptake and glutaminolysis in Th17 cells to reduce IL-17A-mediated inflammation, which could have broad implications for future studies in immunometabolism as well as therapeutics.

## RESULTS

### Metabolic markers in human CD4+ T cells from the mediastinal lymph nodes are decreased in men compared to women.

Metabolic markers are differentially expressed based on the metabolic requirements of CD4+ T cells (Ahl et al., 2020; Healey *et al*., 2021; Kono *et al*., 2018). Using excised mediastinal lymph nodes from deceased human donor males and females that were matched for age, race, and ethnicity (Table S1), we conducted mass cytometry (CYTOF) using an antibody panel focused on T cells and metabolic markers (Table S2). Surface marker expression was used to identify various CD4+ T cells subsets, and there were no significant differences in T and B cell populations (Figure 1A-B). Mean expression of metabolic markers were also determined in CD4+ T cell subsets. Metabolic markers included Glut1 for glycolysis, GLUD1 for glutaminolysis, Cpt1a for fatty acid metabolism, and Grim19, ATP5a, and cytochrome c as various parts of the electron transport chain in mitochondrial metabolism. Th2 and Th17 cells from males had decreased expression of GLUD1, Grim19, ATP5a, and cytochrome c when compared to Th2 and Th17 cells from females (Figure 1C-F). Further, GLUD1 and cytochrome c expression were decreased in Th1 cells from males compared to females (Figure S1A), and GLUD1, ATP5a, and Grim19 expression were decreased in Tregs from males compared to females (Figure S1B).

**Figure 1.**
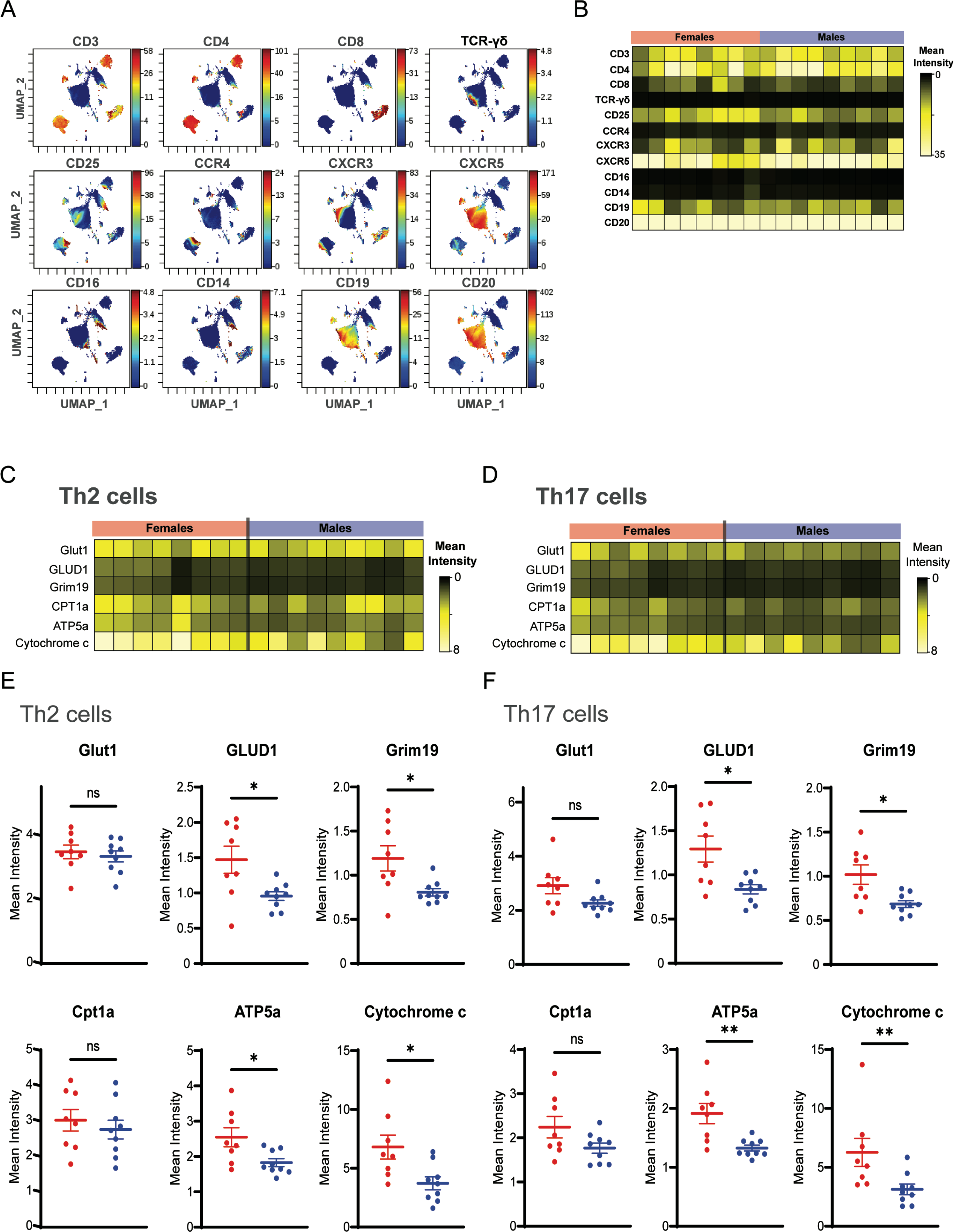
CYTOF of lung draining lymph nodes of deceased donor patients reveals sex differences in T cells and metabolic protein expression. CYTOF was conducted on human lung draining lymph nodes from de-identified female (n=8) and male (n=9) deceased donors. A. UMAP visualization of cell surface markers in all samples. B. Mean expression of surface markers in males and females. C. Mean expression of metabolic markers on Th2 cells (gated on Live, CD3+, CD4+, CXCR5-, CXCR3-, CD25-, CCR4+). D. Mean expression of metabolic markers on Th17 cells (gated on Live, CD3+CD4+, CXCR5-, CXCR3-, CD25-, CCR4-). E-F. Quantification of mean expression data from C and D (mean ± SEM). *p<0.05, **p<0.01, ns: not significant, two-tailed Mann-Whitney U test. See Figure S1 and Tables S1-S2.

These data show that males have decreased expression of glutamine and oxidative phosphorylation-related enzymes in effector CD4+ T cells from human lung draining lymph nodes, suggesting numerous sex-specific differences in CD4+ T cell metabolic dependencies.

### Androgen receptor signaling intrinsically decreases Th17- but not Th2-mediated inflammation

Based on this sex difference in Th2 and Th17 metabolic expression and our previous reports that androgen receptor (AR) signaling decreases type 2 and non-type 2 inflammation during allergic airway inflammation (Cephus et al., 2017; Fuseini *et al*., 2018; Gandhi *et al*., 2022; Kalidhindi et al., 2021), we next determined if AR signaling directly on CD4+ T cells attenuated house dust mite (HDM)-induced airway inflammation. *Cd4^Cre^ Ar^fl/0^* male and *Ar*^fl/0^ male mice were challenged with HDM (40μg) 3 times a week for 3 weeks, and lungs and bronchoalveolar lavage (BAL) fluid were harvested one day after the last challenge to assess airway inflammation (Figure 2A). HDM-challenged *Cd4^Cre^ Ar^fl/0^* male mice had increased neutrophil infiltration and no difference in eosinophils in the BAL fluid compared to HDM challenged *Ar^fl/0^* male mice (Figure 2B). Further, HDM-challenged *Cd4^Cre^ Ar^fl/0^* male mice had increased numbers of lung Th17 cells compared to HDM challenged *Ar^fl/0^* male mice (Figure 2C-D, Figure S2A). No differences in lung Th2 cells were noted between *Cd4^Cre^ Ar^fl/0^* males and

**Figure 2.**
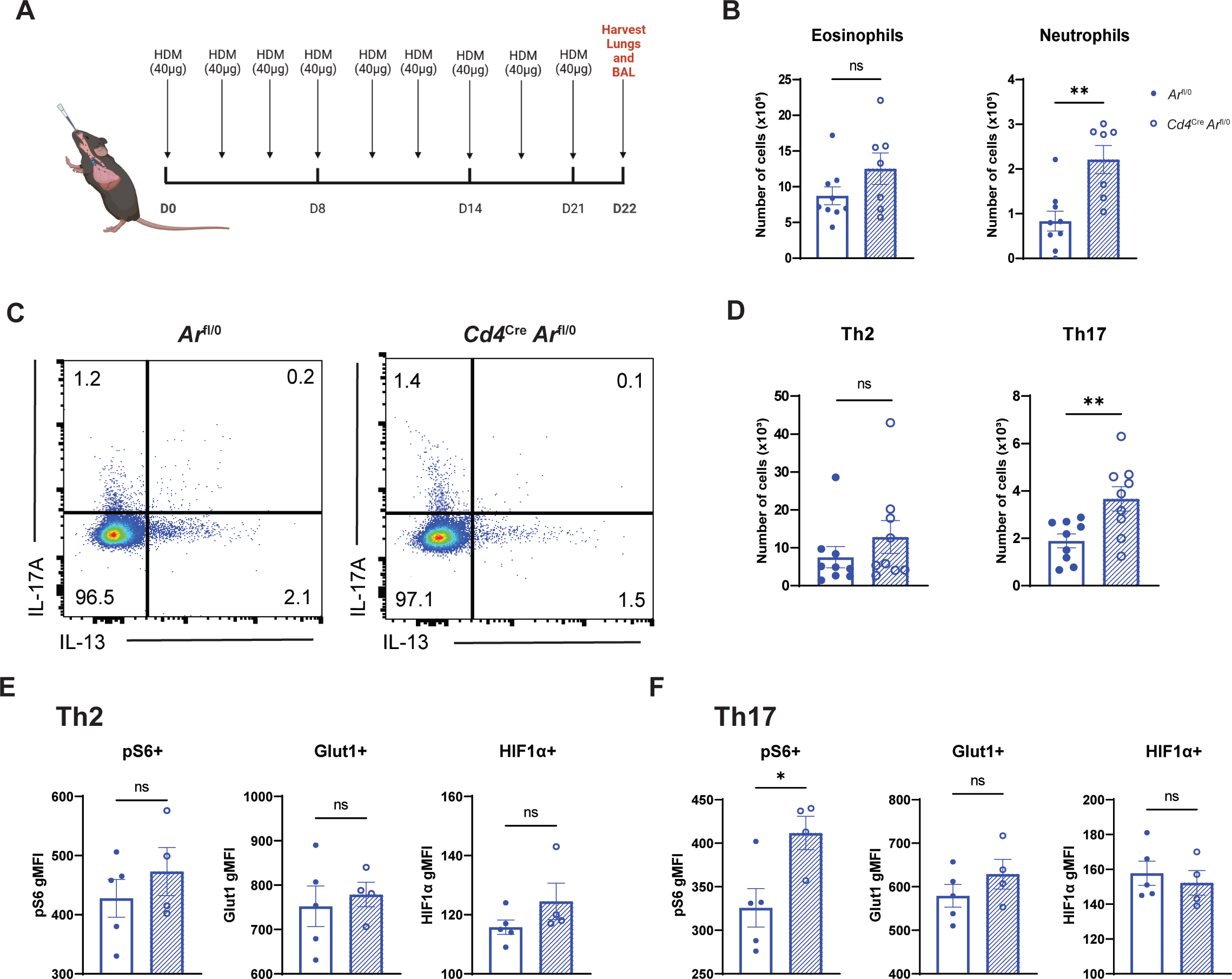
AR signaling in CD4+ T cells reduces Th17 driven neutrophilic inflammation during airway inflammation. A. Model of house dust mite (HDM) allergen challenge. Male *Ar^fl/0^ and Cd4^Cre^ Ar^fl/0^* mice were challenged intranasally with 40µg of HDM three times per week for three weeks. **B.** Quantification of eosinophils and neutrophils in the BAL fluid from mice after HDM challenge (n=8-9 mice per group, BAL could not be collected on one mouse). **C.** Representative flow diagrams of IL-13+ Th2 cells and IL-17A+ Th17 cells. **D.** Quantification of Th2 and Th17 cells in the lung after HDM challenge (n=8-9 mice per group). **E-F.** Expression of pS6 HIF1α, GLUD1, and Glut1 in lung Th2 (**E**) and Th17 (**F**) cells after HDM challenge (n=4-5 mice per group, representative of 2 experiments). All graphs show mean ± SEM, *p<0.05, **p<0.01, ns: not significant unpaired T test. See Figure S2.

*Ar*^fl/0^ male mice (Figure 2C-D). These data were expected since our prior studies using an *in vivo* adoptive transfer model showed AR signaling had no direct effect on Th2 cells (Fuseini *et al*., 2018). In parallel experiments, we also examined if AR signaling in CD4+ T cells impacted HDM-induced airway inflammation in female mice using the *Cd4^Cre^ Ar^fl/fl^* and *Ar^fl/fl^* female mice.

No significant differences in eosinophils or neutrophils in the BAL fluid nor differences in lung Th2 and Th17 cells were noted between the HDM challenged *Cd4^Cre^ Ar^fl/fl^* and *Ar^fl/fl^* female mice (Figure S2B-C).

Using flow cytometry, we also determined expression levels of the metabolic markers pS6, Glut1, and HIF1α in Th2 and Th17 cells. pS6 is a marker for mTORC activation, a key regulator in T cell activation and metabolism. HIF1α is a key metabolic regulator for Th17 cells (Shi et al., 2011). In Th2 cells, no differences were noted in metabolic enzyme expression regardless of AR expression on CD4+ cells (Figure 2E). pS6 expression was increased in Th17 cells from *Cd4^Cre^ Ar^fl/fl^* from male mice compared to cells from *Ar^fl/fl^* male mice (Figure 2F). There were no significant differences in metabolic enzyme expression in Th2 or Th17 cells from *Cd4^Cre^*

*Ar^fl/fl^* and *Ar^fl/fl^* females (Figure S2D-E). This was not surprising given the low levels of circulating androgen in females. Together, these data suggest that AR signaling intrinsically decreased HDM-induced, Th17-mediated inflammation through decreases in metabolic function as indicated by decreased pS6 expression.

### AR signaling reduces mitochondrial metabolism in differentiated Th17 cells

Given our findings that AR signaling decreased metabolic markers in CD4+ cells in humans and Th17 cells in mice, we next determined if AR signaling functionally decreases Th17 and Th2 metabolism. Naïve splenic CD4+ T cells from wild-type (WT) male and *Ar*^Tfm^ male mice were differentiated into Th2 and Th17 cells. *Ar*^Tfm^ mice have a global mutation in the androgen receptor making it nonfunctional, and we confirmed our previous findings that Th17 cells from *Ar*^Tfm^ male mice have increased expression of *Il17a* compared to WT males after 3 days of *in vitro* differentiation (data not shown) (Fuseini *et al*., 2018). We next determined mitochondrial respiration and glycolysis using extracellular flux analysis (van der Windt et al., 2016). Th17 cells from *Ar*^Tfm^ male mice had increased basal respiration, maximal respiration, and ATP production, and trended towards increased spare respiratory capacity compared to Th17 cells from WT mice (Figure 3A-B). Consistent our *in vivo* allergen model, AR signaling had no direct effects on Th2 mitochondrial respiration *in vitro* (Figure S3A-B). AR signaling did not significantly impact basal glycolysis, glycolytic capacity, or glycolytic reserve in Th17 or Th2 cells (Figure 3C-D and Figure S3C-D, respectively).

**Figure 3.**
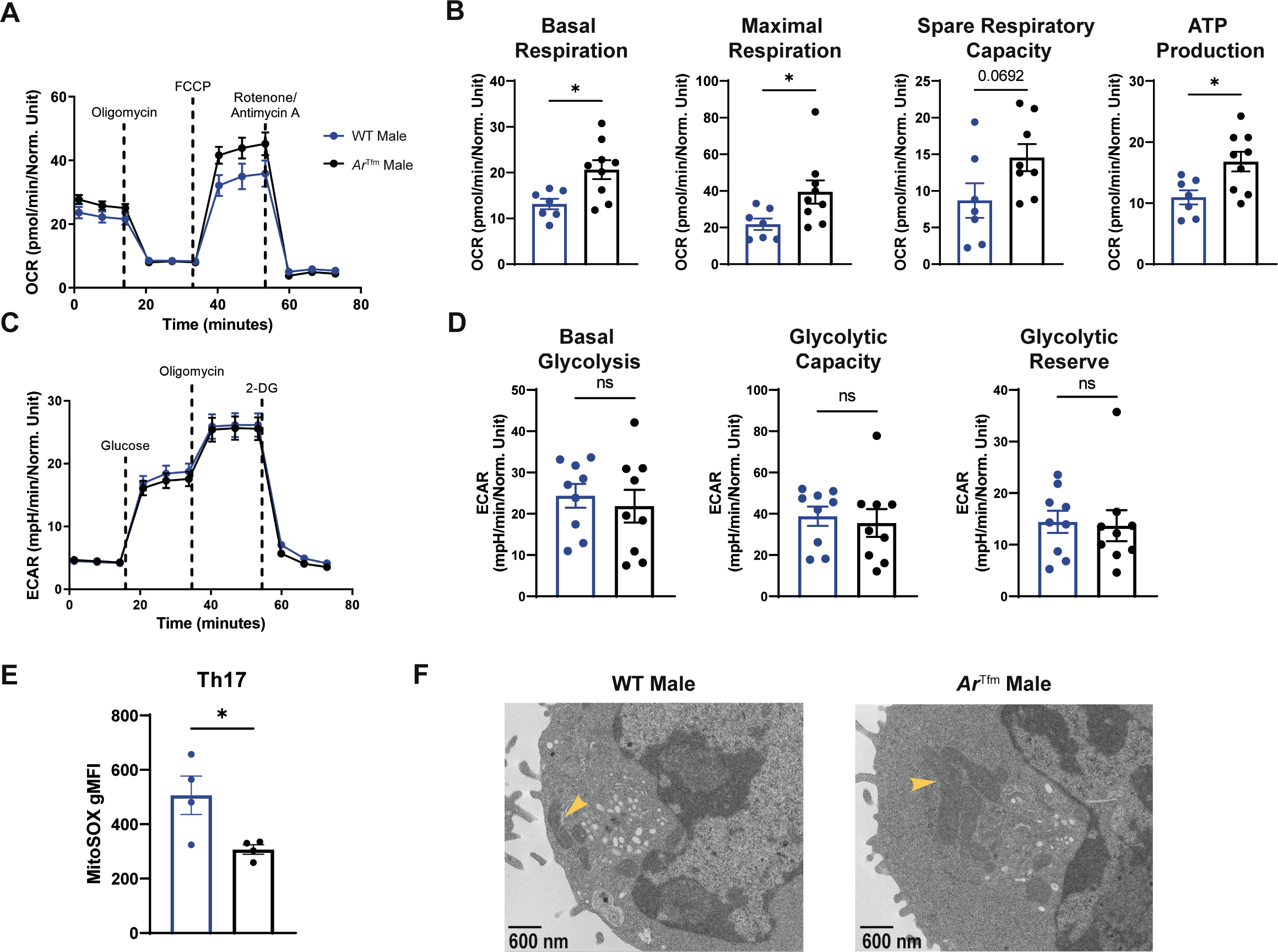
AR signaling modifies mitochondrial metabolism but not glycolysis in Th17 cells. A. Seahorse MitoStress Test on differentiated Th17 cells from WT male and *Ar^Tfm^* male mice measures mitochondrial respiration using oxygen consumption rate (OCR) (n=7-8 mice per group). **B.** Quantified measures of basal respiration, maximal respiration, spare respiratory capacity, and ATP production from panel A. **C.** Seahorse GlycoStress test on differentiated Th17 cells from wild-type male and *Ar^Tfm^* male mice to measure glycolysis using extracellular acidification rate (ECAR) (n=9 mice per group). **D.** Quantified measures of basal glycolysis, glycolytic capacity, and glycolytic reserve from panel C. **E.** Expression of MitoSOX Red, a mitochondrial superoxide marker, in differentiated Th17 cells from WT male and *Ar^Tfm^* male mice (n=4 mice per group, representative of 2 independent experiments). **F.** Representative transmission electron microscopy of mitochondria from differentiated Th17 cells from WT male and *Ar^Tfm^* male mice (yellow arrow indicates a mitochondrion). All graphs show mean ± SEM, *p<0.05 or as shown, ns not significant, unpaired T test. See Figure S3.

To further characterize the impact of AR signaling on Th17 and Th2 cell mitochondria and understand whether AR signaling modified the generation of reactive oxygen species (ROS), we measured MitoSox Red via flow cytometry. ROS suppress Th17 differentiation and effector function (Gerriets *et al*., 2015; Kaufmann et al., 2019; Mak et al., 2017). Consistently, Th17 cells from *Ar*^Tfm^ male mice had decreased mitochondrial superoxide production compared to Th17 cells from WT male mice (Figure 3E), but AR signaling had no effect on ROS production in Th2 cells (Figure S3E). Next, we determined if AR signaling modified mitochondrial morphology by electron microscopy (EM). Mitochondria from *Ar*^Tfm^ male mice were larger and more numerous compared to mitochondria from WT male mice, showing that AR signaling decreased mitochondrial size and number which has been associated with decreased mitochondrial respiration and Th17 effector function (Figure 3F) (Baixauli *et al*., 2022; Buck *et al*., 2016; Hong *et al*., 2022). Collectively, these data show AR signaling specifically restricts oxidative metabolic output in Th17 cells, though the exact mechanisms and pathways remained unclear.

### AR signaling reduces glutamine metabolism in differentiated Th17 cells

Since AR signaling decreased mitochondrial metabolism in Th17 cells, we next conducted targeted metabolomics and pathway analysis on differentiated Th17 cells from WT male and *Ar*^Tfm^ male mice (Pang et al., 2021). Glutamine metabolic pathways were significantly increased in Th17 cells from *Ar*^Tfm^ male mice compared to Th17 cells from WT males (Figure 4A, Table S3). Th17 cells from *Ar*^Tfm^ male mice also had increased intracellular glutamate and a trend for increased glutamine compared to Th17 cells from WT males (Figure 4B).

**Figure 4.**
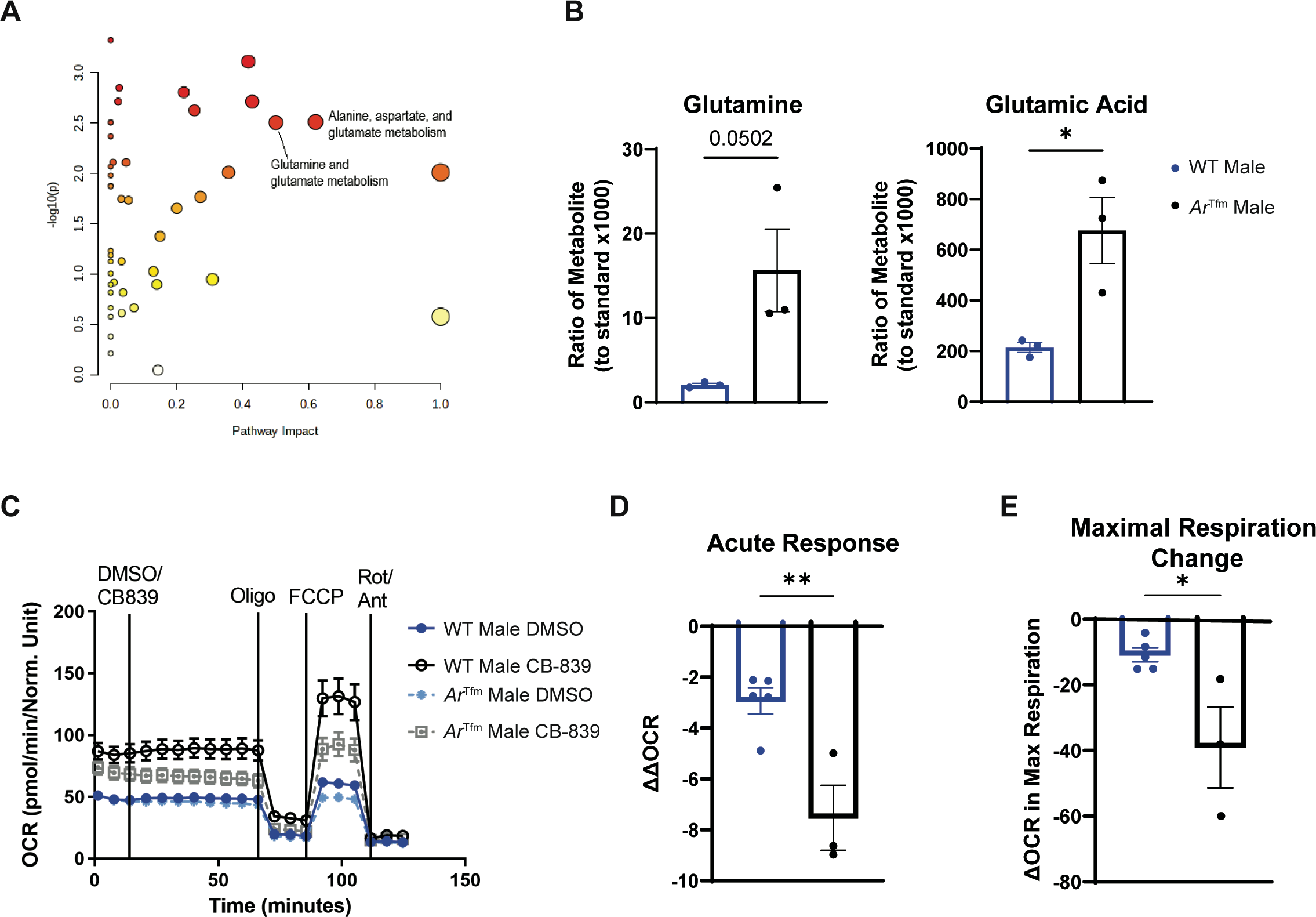
AR signaling reduces glutamine metabolism in Th17 cells. A. Pathway Analysis on targeted metabolomics from differentiated Th17 cells from wild-type male and *Ar^Tfm^* male mice (n=3 mice per group, statistical analysis by MetaboAnalyst Global test, larger circle indicates larger impact, darker shade indicates increased significance). **B.** Intracellular glutamine and glutamate levels from differentiated Th17 cells from WT male and *Ar^Tfm^* male mice measured by mass spectrometry (n=3 mice per group). **C.** Seahorse Substrate Oxidation Assay using CB-839, an inhibitor of glutaminase, on differentiated Th17 cells from wild-type male and *Ar^Tfm^* male mice to measure dependance of mitochondrial respiration of Th17 cells on glutamine metabolism. **D.** Quantified ΔΔOCR from C. Δ difference in measured OCR after DMSO or CB-839 injection. ΔΔOCR was calculated as follows: Δ – Δ . **E.** Quantified from C and calculated as OCR – OCR_DMSO Max_. *p<0.05, **p<0.01, or p-value as shown, unpaired T test unless otherwise specified. See Figure S4 and Table S3.

We next wanted to determine the contribution of glutaminolysis to oxidative phosphorylation in Th17 cells from WT and *Ar*^Tfm^ male mice by conducting a Seahorse Substrate Oxidation Assay using CB-839, an inhibitor of GLS (Voss et al., 2021). To conduct this assay, basal respiration was determined during the first 10 minutes of the assay. CB-839 (10 μM) or vehicle control (DMSO) was then injected onto Th17 cells (at 10 minutes) and incubated for 60 minutes prior to starting a standard Seahorse MitoStress assay (Figure 3C). As shown graphically in Figure S4, ΔOCR_basal_ was determined by subtracting the basal OCR measurement at the end of the CB839 incubation period from the basal OCR at the time of CB- 839 or vehicle injection. ΔΔOCR_basal_ was calculated as ΔOCR_CB-839 basal_ – ΔOCR_DMSO basal_, providing a measure of how reliant the cells were on glutaminolysis (Figure S4). Th17 cells from *Ar*^Tfm^ male mice had a greater decrease in ÄÄOCR_basal_ and decreased ⊗OCR_max_ compared to Th17 cells from WT male mice (Figure 4D-E). These results show that AR signaling decreases glutaminolysis-dependent oxidative phosphorylation in Th17 cells, providing a mechanism for AR-dependent decreases in Th17 differentiation and function.

### Disruption of glutaminolysis alleviates airway inflammation in female but not male mice

We further explored if there was a sex difference in the reliance of glutaminolysis in lung Th17 cells *in vivo*. *Cd4^Cre^ Gls^fl/fl^* and *Gls^fl/fl^* male and female mice underwent the HDM protocol outlined in Figure 2A. Female *Cd4^Cre^ Gls^fl/fl^* mice had decreased IL-13 and IL-17A production in whole lung homogenates, decreased eosinophils and neutrophils in the BAL fluid, and decreased lung Th2 and Th17 cells compared to female *Gls^fl/fl^* mice (Figure 5A-D, Figure S5).

**Figure 5.**
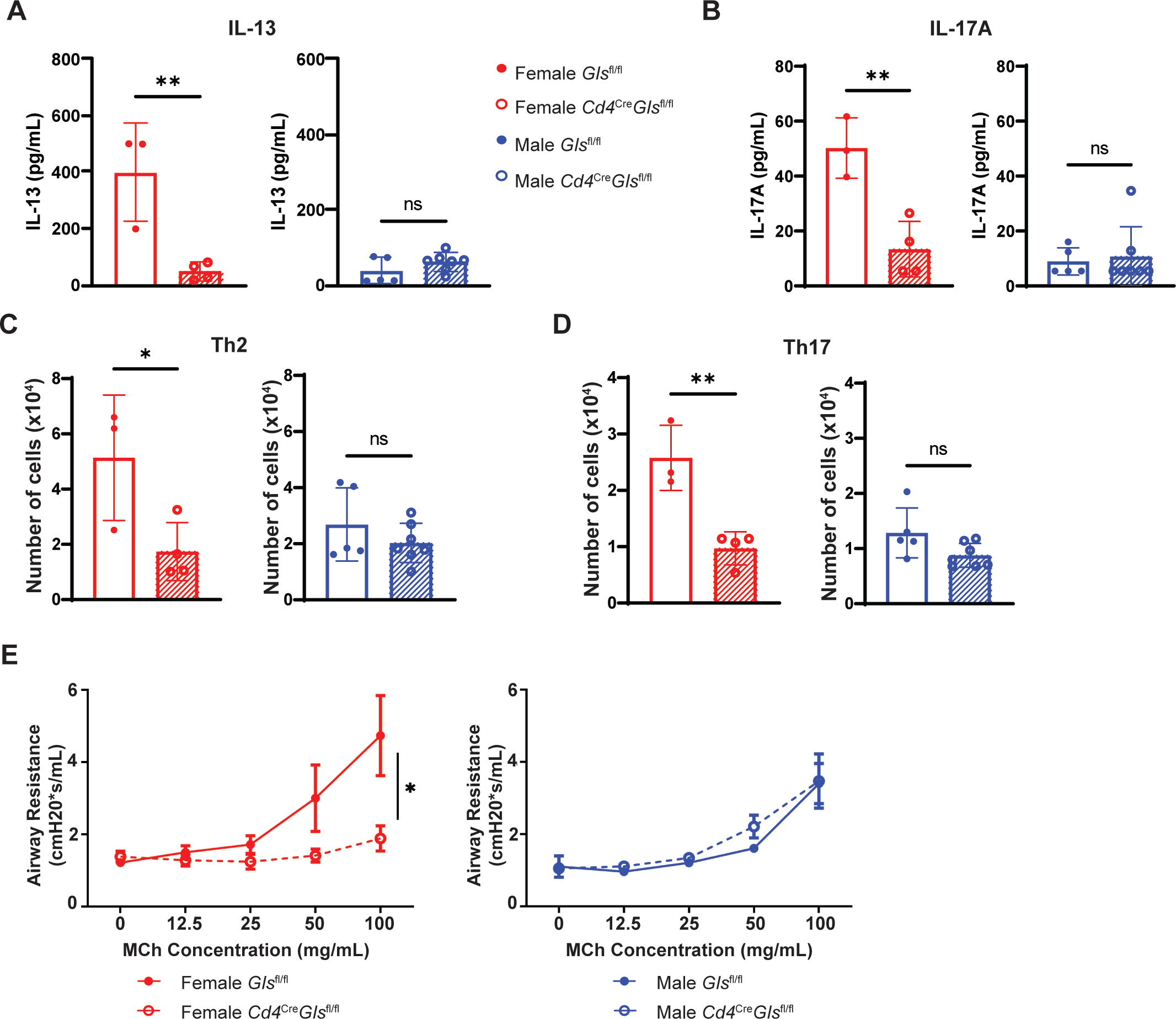
CD4-specific knockout of glutaminase (*Gls*) significantly decreases allergic airway inflammation in female mice but not male mice. A-B. **Total IL-13** (A) **and IL-17A** (B) in lung homogenates measured by ELISA. **C.** Quantification of Th2 cells by flow cytometry. **D.** Quantification of Th17 cells by flow cytometry. **E.** Airway hyperresponsiveness as measured by FlexiVent with increasing doses of methacholine. n=3-6 mice per group, *p<0.05, ** p<0.01, ns not significant. Panels A-D is unpaired T test and panel E is ANOVA of repeated measures with Bonferroni post hoc analysis. Experiments are representative of two independent experiments. See Figure S5.

No differences in any of these endpoints were determined between HDM challenged *Cd4^Cre^ Gls^fl/fl^* and *Gls^fl/fl^* male mice. Further, *Cd4^Cre^ Gls^fl/fl^* female mice had decreased airway hyperresponsiveness to methacholine, a physiological hallmark of asthma, compared to *Gls^fl/fl^* female mice (Figure 5E). This change in airway hyperresponsiveness was not detected in HDM challenged *Cd4^Cre^ Gls^fl/fl^* and *Gls^fl/fl^* male mice. These findings show that Th2 and Th17 cells from males, with high AR signaling, do no rely on glutaminolysis during allergic airway inflammation and support findings from Figures 3-4. Further, these findings show that glutaminolysis in Th2 and Th17 cells is required for maximal HDM-induced airway inflammation, providing an *in vivo* mechanism for increased Th2 and Th17 cells in HDM challenged females compared to males.

### AR signaling reduces glutamine uptake in Th17 cells

Based on our *in vitro* and *in vivo* findings of AR signaling differentially impacting glutaminolysis reliance, we conducted *in vitro* and *in vivo* glutamine metabolism-targeted CRISPR screens using OVA-specific Th17 differentiated cells from OT-II Cas9 male and female mice to test which specific genes in the pathway were responsible for the sex-specific differences in glutamine reliance (Figure S6A, Figure 6A) (Sugiura *et al*., 2022). In the *in vitro* CRISPR screen, a single glutaminolysis-related gene was knocked out in each OVA-specific Th17 cell using guide RNAs (gRNA). After 7 days in culture OT-II Cas9 Th17 cells were recovered and sequenced for their gRNA abundance. Depletion of a gRNA (a negative log fold change) indicates that gene is essential for Th17 fitness and proliferation. Enrichment of a gRNA (a positive log fold change) indicates that gene functions as a repressor of proliferation in Th17 cells. As shown in Figure S6B, the *in vitro* CRISPR screen revealed sex differences in metabolic genes that impacted Th17 proliferation with significant differences in females denoted in red, significant differences in males denoted in blue, and significant differences in both males and females denoted in purple. *Slc1a5*, which encodes the glutamine transporter ASCT2, was essential for female Th17 cell fitness *in vitro* (Figure S6B). To support our findings from the CRISPR screen Th17 cells from *Ar^T^*^fm^ male mice had increased *Slc1a5* compared to Th17 cells from male mice (Figure S6C). These findings show a sex difference in glutaminolysis reliance in Th17 cells and that AR signaling decreases the expression of the glutamine transporter ASCT2 (*Slc1a5*).

**Figure 6.**
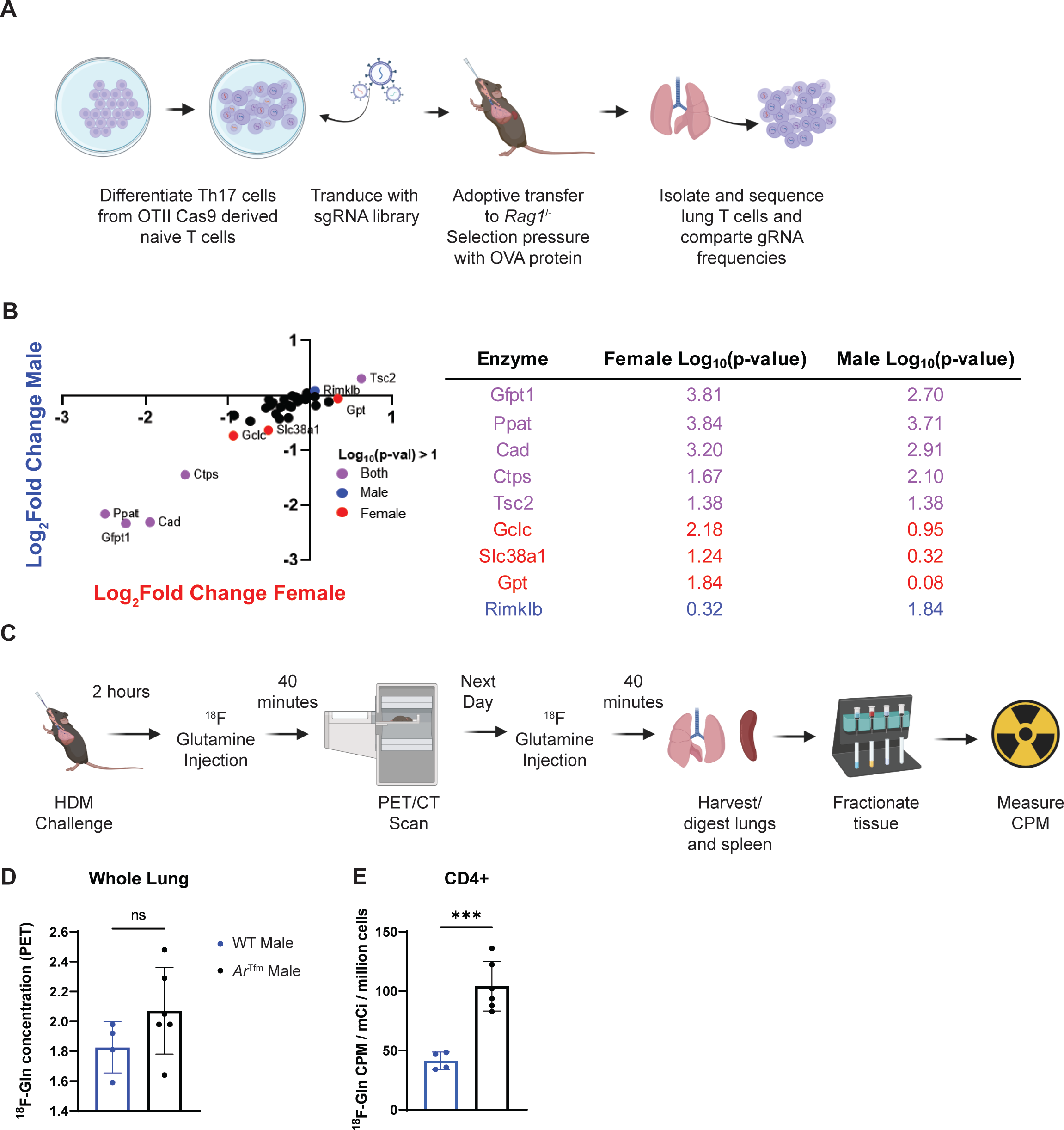
AR signaling reduces glutamine uptake in T cells. A. Model of targeted CRISPR screen using a glutamine library on differentiated Th17 cells from male and female OT-II Cas9 mice in an OVA-induced lung inflammation model. **B.** Change in gRNA abundance in lung Th17 cells from glutamine metabolism targeted CRISPR screen after OVA-induced lung inflammation with table showing statistics (n=4-5 mice, statistical analysis by MAGeCK, significantly affected represented by color of dot as shown in legend). **C.** Model of F18-Glutamine PET studies in HDM-induced lung inflammation (as shown in Figure 2A). **E.** Quantification of ^18^F-Glutamine concentration in whole lung by PET/CT imaging (n=4-6 mice per group). **F.** Quantification of ^18^F- Glutamine concentration in lung CD4+ T cells by magnetic separation and gamma counting, normalized to viable cells (n=4-6 mice per group). ***p<0.001, ****p<0.0001, unpaired T test unless otherwise specified. See Figure S6.

A similar experiment was performed *in vivo* where Th17 differentiated cells from OT-II Cas9 male and female mice were transduced with gRNAs from a glutamine CRISPR library and then adoptively transferred into *Rag1*^-/-^ male mice. *Rag1*^-/-^ recipient mice were administered OVA protein intranasally to establish OVA-induced airway inflammation. Lungs were harvested one day following the last OVA challenge and OVA-specific T cells were isolated by cell sorting and gRNAs sequence. *Slc38a1*, another glutamine transporter SNAT1, was significantly depleted in females and required for Th17 cell fitness (Figure 6B). Collectively, our *in vitro* and *in vivo* CRISPR screens show a sex difference in glutamine uptake genes for Th17 cells.

To directly test if AR signaling decreased glutamine uptake in CD4+ T cells during ongoing allergic airway inflammation, we conducted an *in vivo* ^18^F labelled glutamine uptake assay. As shown in the model in Figure 6C, we used ^18^F-glutamine tracing and magnetic cell separation to determine glutamine uptake in various cell types as previously shown (Reinfeld et al., 2021). ^18^F-glutamine levels were not significantly different in the whole lung by whole body PET/CT scan (Figures 6D, S6D), but glutamine uptake by CD4+ cells in the lungs was increased in *Ar^T^*^fm^ male mice compared to the WT male mice (Figure 6E). Glutamine uptake was not different in CD4+ cells or CD4- cells in *Ar^T^*^fm^ male mice compared to the WT male mice in the spleen (Figure S6E), showing tissue localized changes in CD4+ cell metabolism with intranasal HDM challenge. Interestingly, HDM-challenged *Ar^T^*^fm^ male mice also had increased glutamine uptake in other cells, including CD11b-CD4- lung cells compared to WT male mice, indicating effects of AR signaling on glutaminolysis in other cell types in the lung (Figure S6F).

In all, AR signaling reduced glutamine uptake in lung CD4+ cells during allergic airway inflammation, supporting a mechanism for decreased reliance of glutaminolysis in male Th17 cells.

### AR signaling reduces Th17 metabolism in human cells

Our murine *in vitro* and *in vivo* data demonstrated that AR signaling decreased expression of glutamine transporters and decreased glutaminolysis in Th17 cells, providing a mechanism for decreased reliance of glutaminolysis in male Th17 cells. We wanted to confirm this mechanism in circulating CD4+ T cells from men and women with severe asthma (Table S4) using a single-cell metabolic approach, SCENITH (Figure 7A)(Argüello et al., 2020).

**Figure 7.**
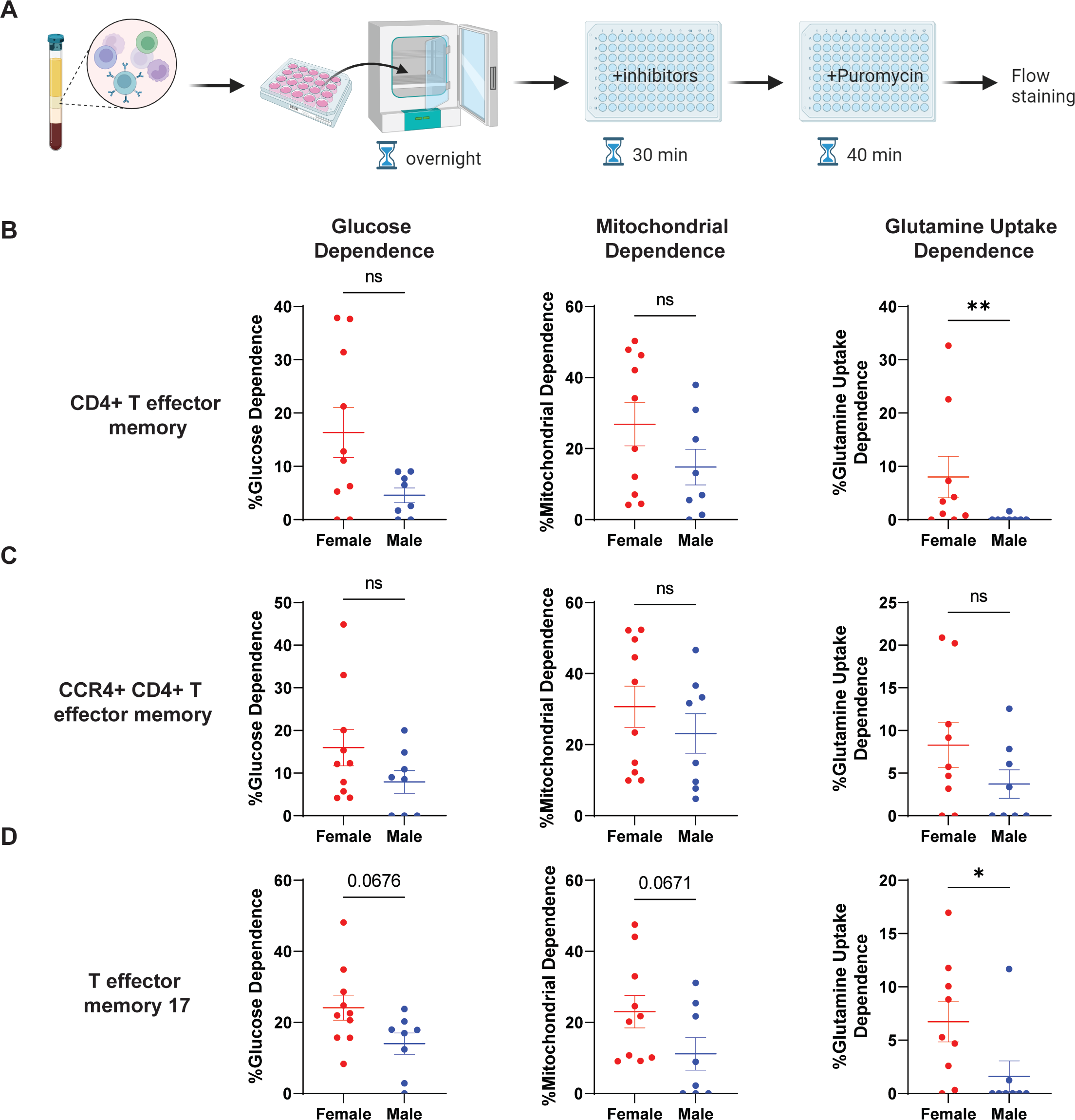
Males with severe asthma have decreased dependence upon glutamine uptake in circulating CD4+ T cell subsets compared to females with severe asthma. A. Model of SCENITH protocol on frozen PBMCs restimulated overnight with antiCD3/CD28/CD2. B-D. Glucose dependence, mitochondrial dependence, and glutamine uptake dependence in males and females measured using SCENITH inhibitors (2-deoxyglucose, oligomycin, and V9302, respectively) in (A) CD4+ T effector memory cells (defined as Live, CD3+, CD4+, CD8-, CD45RA-, CCR7-), (B), CCR4+ CD4+ T effector memory cells, and (C) T effector memory 17 cells (defined as B and CCR4+, CCR6+). *p<0.05, **p<0.01, ns: not significant, or p-value as shown; Mann-Whitney U test. See Figure S7.

SCENITH is a flow-based single cell metabolic technique that relies on puromycin to track protein synthesis in the presence of metabolic pathway inhibitors – 2-deoxyglucose (2-DG) inhibiting glycolysis, oligomycin inhibiting mitochondrial respiration, and V9302 (Schulte et al., 2018) inhibiting glutamine uptake to calculate the dependence of specific cells on these metabolic pathways. There were no significant differences between male and female CD4+ T cells in glucose dependence or mitochondrial dependence, but there was a significantly decreased dependence on glutamine uptake from male CD4+ T cells compared to female CD4+ T cells (Figure S7A). Circulating CD4+ T effector memory cells (TEMs) had significantly decreased dependence on glutamine uptake via ASCT2 in male cells compared to female cell but there were no sex differences in glucose or mitochondrial dependence (Figure 7B). CCR4+ TEMs, encompassing Th2 and Th17 cells, had no significant differences in glucose, mitochondrial, or glutamine uptake dependence between male and female cells (Figure 7C).

Finally, circulating Th17 cells, defined as CCR4+CCR6+ TEMs, from males exhibited a trend towards decreased dependence on glucose and mitochondrial metabolism and a significant difference in glutamine uptake dependence for metabolism compared to Th17 cells from females (Figure 7D). Other CD4+ T cells were examined, including naïve CD4+ T cells and T central memory cells (TCMs). Naïve CD4+ T cells from males had significantly decreased reliance upon glutamine uptake but no difference in glucose or mitochondrial dependences compared to naive CD4+ cells from females (Figure S7B). In CD4+ TCMs, males had a trend towards decreased reliance upon glucose and significantly decreased reliance upon glutamine uptake (Figure S7C). There were no differences by sex in mitochondrial dependence in CD4+ TCMs. These data confirm that in human Th17 cells, but not Th2 cells, AR signaling decreases reliance on glutamine uptake.

Our mouse and human data show that male sex and AR signaling decrease glutamine uptake and/or glutaminolysis in Th17 cells, yet it is still unclear how AR signaling decreases expression of glutamine transporters and enzymes. As our culture systems and ex vivo human experiments, which had similar amounts of androgens in the media, demonstrated sex differences and the effects of AR on Th17 cells, and we observed sex differences in glutamine uptake-dependence even in naïve CD4+ T cells, we explored whether epigenetic changes could play a role. Prior studies showed that trimethylation at histone 3 lysine 27 (H3K27me3) is associated with repressor function and H3K27 demethylase enzymes are critical in maintaining metabolic genes necessary for Th17 function (Cribbs et al., 2020; Li et al., 2014a; Wei et al., 2009). Therefore, we explored if there were differences H3K27me3 signatures at sites related to glutamine uptake and glutaminolysis using the publicly available ENCODE data (https://www.encodeproject.org/) with CHIP-sequencing data from 2 males (identifiers: ENCFF993RMD, ENCFF820GJE) and 1 female (identifier: ENCFF984ZEE) (Consortium, 2012; Luo et al., 2020). We determined that circulating T cells from 2 males had increased H3K27Me3 marks at the *Slc1a5* locus compared to the female sample, though we lacked sufficient samples to determine statistical significance (Figures S7D). Altogether, these data indicate that AR signaling reduces dependence on glutamine uptake in CD4+ T cells, and specifically in Th17 cells, similar to our findings in mice, indicating a sex-specific regulation of nutrient uptake to limit inflammation.

## DISCUSSION

There is a female predominance in lupus, rheumatoid arthritis, and severe asthma (Chowdhury *et al*., 2021; Klein and Flanagan, 2016). CD4+ T cells are important drivers of inflammation in these diseases, yet, how sex hormones directly modify CD4+ T cell effector responses remains unclear. In this study, we employed *in vitro* and *in vivo* methods in mice and human T cells to show that AR signaling restricted Th17 effector function through decreased glutamine uptake and glutaminolysis. Our findings identify potential precision medicine targets by elucidating differential mechanisms of T cell metabolism by sex hormone signaling.

Previous studies by our laboratory and others showed AR signaling decreased Th17- and Th2- mediated inflammation (Cephus *et al*., 2017; Fuseini *et al*., 2018; Gandhi *et al*., 2022; Kalidhindi *et al*., 2021). In this study, we defined a foundational mechanism and showed that AR signaling decreased CD4+ T cell glutamine uptake and subsequent glutaminolysis, that male mice with normal androgen levels and AR signaling had decreased CD4+ T cell dependence on glutaminolysis pathways, and that CD4-specific deletion of *Gls* decreased Th2 and Th17-aiway inflammation only in female mice. Further, we found that AR signaling decreased glutamine uptake transporters, such as ASCT2 or SNAT1. These results provide an intrinsic pathway for how AR signaling attenuates glutaminolysis leading to attenuated Th17 cell differentiation and effector function in males.

Glutaminolysis is important for both Th17 and Th2 effector function (Healey *et al*., 2021; Johnson *et al*., 2018; Kono *et al*., 2018). Importantly, we studied mediastinal lymph nodes collected from deceased donors and performed CYTOF (thus studying a specific tissue rather than peripheral blood) and discovered there was a sex bias in CD4+ T cell metabolic enzyme expression in various subsets. Importantly, we found that GLUD1 expression and expression of multiple proteins in the electron transport chain were significantly decreased in Th2 and Th17 cells in the lymph nodes of men, indicating sex differences in the metabolic processes of these cells. As our panel was limited by the available antibodies and panel size, we acknowledge that other pathways could potentially be differentially regulated as well. Additionally, our analysis was limited to CD4+ T cells, but there are likely many differences in other cell types.

AR signaling decreasing glutamine uptake in CD4+ T cells and subsequent glutaminolysis in Th17 cells is likely a result in AR signaling shunting glutamine metabolism away from the TCA cycle and towards reactive oxygen species management. Increased reactive oxygen species limited the Th17 differentiation and effector function by promoting a regulatory phenotype (Gerriets *et al*., 2015; Johnson *et al*., 2018; Kim et al., 2014; Won et al., 2013). Further, increased methylation, especially at H3K27me3, decreased metabolic function in Th17 differentiation (Cribbs *et al*., 2020; Li *et al*., 2014a). Previous studies showed the accessibility of these genes, especially regarding H3K27me3 via KDM6, is important in the differentiation and maintenance of Th17 effector function (Cribbs *et al*., 2020). This repressor function can impact a multitude of pathways, most important being mitochondrial biogenesis and respiratory activity.

These findings, along with previous studies on the importance of mitochondrial dynamics and function in Th17 cells (Baixauli *et al*., 2022; Buck *et al*., 2016; Hong *et al*., 2022) align with our functional findings regarding decreased mitochondrial respiration and qualitative findings regarding changes in the mitochondria. Therefore, it is possible that AR signaling impacts KDM6 mediated H3K27me3 methylation on multiple Th17-related pathways and this can be explored in future studies. Additional studies are also needed on the effect of AR signaling on acetylation pathways, as previous studies have indicated effects of polyamine metabolism on acetylation of regions related to CD4+ T helper differentiation (Puleston *et al*., 2021).

Nutrient uptake is important in driving T cell differentiation and effector function (Buck *et al*., 2015; Wei *et al*., 2017). While glutamine uptake and utilization became the focus of our study as it was one of our strongest hits, we acknowledge that AR signaling could be driving differential uptake and utilization of other nutrients, as shown in the other significant hits in our metabolomics studies. Arginine and serine are important in Th17 effector programs and play a role in polyamine and carbon metabolism (Sugiura *et al*., 2022; Wagner et al., 2021). The differences in these pathways with AR signaling could further impact the Th17 differentiation and effector program and the balance of T cells and would be exciting possibilities for future studies.

Many studies have indicated that both ovarian hormones and androgens directly impact the proliferation and function of macrophages, dendritic cells, T regulatory cells, CD8+ T cells, and CD4+ T cells (Chowdhury *et al*., 2021; Klein and Flanagan, 2016). Here, we show a dichotomized reliance upon a specific metabolic pathway based on AR signaling. We found that glutaminolysis is important in Th17 cell effector function in females but may not have as large of an impact on Th17 cells from males. This indicates that AR signaling, while decreasing glutaminolysis, may force Th17 cells to rely upon different metabolic pathways. This will be essential in drug development as more diseases develop immunometabolism targeted therapeutics (Makowski et al., 2020; Pålsson-McDermott and O’Neill, 2020). Our study indicates that those drugs may be differentially effective based on sex and the metabolic mechanisms that are driven by sex hormones. Our metabolomics data and CRISPR screens suggest glutaminolysis and other pathways are differentially regulated by sex hormone signaling in Th17 cells. Future studies on immunometabolism and the targeting of immunometabolism for therapeutics should determine the sex-specific effects of the drugs and how that may impact efficacy. Additionally, the focus of this study on androgens supports the use of weak androgens, such as inhaled DHEA or DHEA-S, as possible therapeutics or adjuvant therapy to increase the effectiveness of other therapeutics (DeBoer et al., 2018; Marozkina et al., 2019; Wenzel et al., 2010). Further studies on the impact of sex hormones on specific metabolic pathways can lead to new targets for therapy and increased effectiveness for those therapeutics and further the field of precision medicine.

## Limitations of the study

The CYTOF panel on our human lymph node studies did not provide any functional metabolic data. Also, while we could determine sex differences in our human studies, we could not specifically measure the role of androgen signaling the human studies. Future studies using patients undergoing gender affirming care (Robinson et al., 2022) or patients with androgen insensitivity syndrome could provide this information. Additionally, all human samples were limited in that they were not timed for menstruation cycles and all medications may not have been captured at the time of death or blood draw. However, androgen levels will not fluctuate to the same degree as ovarian hormones. Our metabolomics study used a targeted panel of 108 metabolites, but we may have missed significantly dysregulated pathways that were not on the panel. Finally, while the glutamine uptake experiments were powerful in determining that AR signaling decreased glutamine uptake in CD4+ T cells, due to the radioactivity of the cells, we were unable to sort specifically on Th17 cells.

## Author Contributions

N.U.C, D.C.N., and J.C.R. designed the research. N.U.C., J.Y.C, M.Z.M, M.M.W, C.C., S.N.K., K.S., and V.D.G. performed the research. A.S., M.T.S, and K.V. performed the research and along with M.Z.M. and M.M.W. provided essential technical support. J.C.R, R.S.P, K.N.C, R.D.G, and A.I.S. provided essential expertise, materials, and samples. N.U.C., J.Y.C., M.Z.M., S.N.K., K.S., X.Y., and D.C.N. analyzed data. N.U.C. and D.C.N. wrote the paper with contributions from the other authors.

## Acknowledgements

We thank members of the Dawn Newcomb, R. Stokes Peebles, Jeffrey Rathmell, and W. Kimryn Rathmell laboratories for their constructive input. Additionally, we would like to thank Heather Pua, Meenakshi Madhur, and Alyssa Hasty for their comments and feedback. We thank members of Anne Sperling’s lab, especially Kelly M. Blaine, for their work banking lung lymph node cells. For providing lung lymph nodes for research, we thank the Gift of Hope Organ and Tissue Donor Network (Itasca, IL) as well as the families of organ donors. We would also like to thank Holly Algood for providing mice. For support with the CYTOF analysis of the lymph nodes, we would like to acknowledge the Human Immune Discovery Initiative (HIDI) of the Vanderbilt Center for Immunobiology (VCI). We would also like to thank Bradley Reinfeld for his help in establishing ^18^F-nutrient assays and Mohammaed Noor Tantawy for his assistance in analysis of the PET images. We would also like to thank H. Charles Manning for providing V9302. We would like to acknowledge the ENCODE Consortium and the lab of Bradley Bernstein at the Broad Institute for the MINT-CHIP-Seq data in the ENCODE database. We acknowledge BioRender and Adobe Illustrator for providing platforms to make figures. Finally, we would like to thank all of patients, donors, participants, and families for their contributions to our project and their commitment to scientific progress. The following cores provided technical assistance on the project: VUMC Flow Cytometry Shared Resource (particularly David Flaherty), Vanderbilt Cancer and Immunology Core (CIC, particularly Caroline Roe and Jonathan Irish), Vanderbilt Cell Imaging Shared Resource (CISR, particularly Evan Krystofiak), Vanderbilt Mass Spectrometry Core Lab (particularly Sergei Chetyrkin), and VANTAGE.

Funding for this project includes F30 HL159941 (N.U.C), T32 GM007347 (N.U.C., M.Z.M, A.S.), F30 CA239367 (M.Z.M), F31 CA261049 (M.M.W.), T32 DK101003 (K.V.), R01 HL136664 (D.C.N, J.C.R), and R01 HL122554 (D.C.N).

## Declaration of Interests

J.C.R. is a founder, scientific advisory board member, and stockholder of Sitryx Therapeutics; a scientific advisory board member and stockholder of Caribou Biosciences; a member of the scientific advisory board of Nirogy Therapeutics; has consulted for Merck, Pfizer, and Mitobridge within the past 3 years; and has received research support from Incyte Corp., Calithera Biosciences, and Tempest Therapeutics.

## Experimental model and subject details

### Human samples

Liquid nitrogen preserved single cell suspensions of de-identified deceased donor derived lymph nodes were provided by Dr. Anne Sperling and Dr. Robert Guzy through their IRB-approved biobank. All lymph nodes were draining lymph nodes from the lung. Patient demographic information is provided in Supplemental table 1 and as indicated in the text.

Peripheral blood mononuclear cells (PBMCs) were collected and banked through an approved IRB (VUMC IRBs122554 and 202162) from patients with severe asthma as defined by the American Thoracic Society criteria (Chung et al., 2014) at the Vanderbilt Asthma, Sinus, and Allergy Program. Patients were 18-45 years old and were excluded if they had any symptoms of infection within the past week, comorbid Th17-associated disease (e.g. Crohn’s disease), or administration of systemic exogenous hormones (e.g. oral contraceptives, estrogen replacement therapy). Women were excluded if they were pregnant, breastfeeding, menopausal, or had undergone an oophorectomy or hysterectomy. Patient demographic information is provided in Supplemental table 2 and as indicated in the text. PBMCs were isolated from whole blood using SepMate-50 PBMC Isolation Tubes (STEMCELL Technologies) and Lymphoprep (STEMCELL Technologies) density gradient isolation. A red blood cell lysis (BioLegend) was performed according to manufacturer instructions. Cells were aliquoted at 10 million cells/mL in BAMBANKER serum-free cell freezing medium (Bulldog Bio) and preserved in liquid nitrogen.

### Mice

All experiments were performed at Vanderbilt University in accordance with Institutional Animal Care and Utilization Committee (IACUC)-approved protocols and conformed to all relevant regulatory standards. Mice were housed in a pathogen-free facility with five mice per cage and ad libitum access to food and water. C57BL/6J, C57BL/6J *Ar^Tfm^*, C57BL6/J *Ar^fl/fl^*, and C57BL6/J *Rag1^-/-^* mice were obtained from Jackson Laboratories and bred in house. OT-II Cas9 double transgenic mice, *Gls^fl/fl^*, and *Cd4^Cre^ Gls^fl/fl^* were provided by the lab of Dr. Jeffrey Rathmell at Vanderbilt University. *Cd4^Cre^* mice that were crossed to *Ar^fl/fl^* mice were provided by the lab of Dr. Holly Algood at Vanderbilt University Medical Center. Mice were genotyped for floxed and Cre alleles, OTII and Cas9 transgenes, or *Ar^Tfm^* or wild-type *Ar* transgenes. Male and female mice were used as described in the text. Eight to seventeen-week-old mice were used for all animal experiments. All mice were considered treatment naïve until the start of the study.

### Cell lines

Plat-E retroviral packaging cell line was provided by the lab of Dr. Jeffrey Rathmell. Cell lines were maintained at 37°C with 5% CO_2_ in DMEM media supplemented with 10% FBS, 100 U/mL penicillin/streptomycin, 1 µg/mL puromycin, and 10 µg/mL blasticidin to maintain expression of viral packaging genes.

### CYTOF on human lymph nodes

CYTOF was run by the Vanderbilt University Cancer and Immunology Core (CIC). Cells were rapidly thawed and recovered in RPMI1640 with L-glutamine (Gibco), 2% FBS, 10mM HEPES, and 100U/mL DNase. Cell labeling and mass cytometry analysis were performed as previously described using the panel located in Table S2 (Kramer et al., 2022). Cells were resuspended in 1X EQ Four Element Calibration Beads and collected on a Helios mass cytometer (Fluidigm) at the Vanderbilt Flow Cytometry Shared Resource Center. CYTOF data was collected, normalized (Finck et al., 2013), and scaled appropriately using Cytobank. Manual gating was used to remove atypical events and a UMAP analysis was performed on cell surface markers to identify immune cell populations. T cell subsets were identified using surface markers and metabolic enzyme expression was quantified using mean expression analysis.

### *In vivo* allergen challenge model

Adult mice were anesthetized with isoflurane and challenged intranasally with 40 µg/75 µL PBS of house dust mite (HDM, Greer) three times per week for three weeks (21 days). Twenty-four hours after the last challenge, mice were sacrificed using 200 µL of pentobarbital sodium for endpoint analysis. In the *in vivo* CRISPR screen, mice were challenged every other day with 25 µg/75 µL PBS of OVA protein starting on day -1 (prior to adoptive transfer of transduced T cells) and ending on day 7. Twenty-four hours after the last challenge, mice were sacrificed for collection of lungs and downstream analysis.

### Bronchoalveolar lavage (BAL) collection

After sacrifice, BAL was performed through the insertion of a tracheostomy tube, instillation of 800 µL of PBS through a syringe into the lungs, gentle massage of the lungs, and gentle withdrawal of fluid through the same syringe. Cells were adhered to a slide using a Cytospin (Thermo Fisher Scientific) and stained using a commercially available Three-Step Stain kit (Richard-Allen Science, Thermo Fisher Scientific). Using light microscopy, eosinophils, neutrophils, macrophages/monocytes and lymphocytes were counted as previously described (Fuseini *et al*., 2018).

### Flow cytometry

Lungs were harvested, minced, and digested using 1 mg/mL collagenase type IV (Sigma-Aldrich) and DNase I (Sigma-Aldrich) in RPMI with 10% FBS (Gibco) for 30 minutes and 37°C (Fuseini *et al*., 2018). Digestion was stopped using 1 µM EDTA. Cells were filtered through a 70 µm strainer to remove debris and a red blood cell lysis was performed per manufacturer instructions (BioLegend). For some experiments, cells were restimulated in IMDM with GlutaMax and HEPES (Gibco), 10% FBS, 1% penicillin/streptomycin, 1% sodium pyruvate, 50 µM 2-mercaptoethanol (Sigma-Aldrich), 1 µM ionomycin (Sigma-Aldrich), 50 ng/mL PMA (Sigma-Aldrich), and 0.07% Golgi Stop (BD Biosciences) at 37°C for 4 hours. Cells were then washed with PBS and 5 million viable cells were used for downstream assays. Cells were stained with a fixable viability dye (Live Dead Aqua, Thermo Fisher Scientific) and blocked using an anti-mouse FcR antibody (anti-CD16/CD32, BD Biosciences) at 4°C. Cells were washed with PBS with 3% FBS and stained for surface markers as stated in the text for 45 minutes at 4°C. Cells were again washed with PBS with 3% FBS and were subsequently fixed and permeabilized using the FoxP3 Transcription Factor Fix/Perm Kit (Thermo Fisher Scientific) per manufacturer instructions for unstimulated cells and Cytofix/Cytoperm Kit (BD Biosciences) for restimulated cells. After wash with the provided Perm Wash (Thermo Fisher Scientific or BD Biosciences), cells were stained for intracellular markers as stated in the text for 45 minutes at 4°C. Cells were washed again with Perm Wash and once more with PBS with 3% FBS. Flow cytometry was conducted on a Cytek Aurora (Cytek Biosciences) and data were analyzed using FlowJo (BD Biosciences).

For detection of mitochondrial superoxide in differentiated Th17 and Th2 cells from mice, cells were washed and stained with surface markers and MitoSox Red (2.5 µM, Thermo Fisher Scientific). Flow cytometry was conducted on a Cytek Aurora and data were analyzed using FlowJo (BD Biosciences).

### ELISAs

Left lungs were flash frozen in liquid nitrogen and homogenized using a bead beater. Cytokine levels were measured using IL-13 and IL-17A Quantikine kits (R&D) per manufacturer’s instructions. Any value below the limit of detection was assigned half the value of the lowest detectable standard.

### Airway hyperresponsiveness

Mice were anesthetized with pentobarbital sodium (85 mg/kg). Using an 18-gauge tracheostomy tube, mice were ventilated using the Sci-Req FlexiVent machine set at 150 breaths/min and tidal volume of 2 mL. Airway responsiveness to methacholine was measured by determining airway resistance (Rrs) at baseline and after successive doses of nebulized-ß-methacholine (0-100 mg/mL; Sigma-Aldrich).

### *In vitro* CD4+ T cell differentiation

Spleens were harvested from appropriate mice, mechanically digested to a single cell suspension, and put through a red blood cell lysis (BioLegend) per manufacturer’s instructions. Primary murine naïve CD4+ T cells were isolated from spleens of mice using the STEMCELL Mouse Naïve CD4+ T Isolation Kit (STEMCELL Technologies) per manufacturer’s instructions. Isolated naïve T cells were plated at 500,000 cells per well in a 24-well non-tissue culture plate coated with anti-CD3 (1 µg/mL, BD Biosciences) and anti-CD28 (0.5 µg/mL, BD Biosciences). Cells were cultured at 37°C with 5% CO_2_ in IMDM with GlutaMax and HEPES (Gibco), 10% FBS, 1% penicillin/streptomycin, 1% sodium pyruvate, and 50 µM 2-mercaptoethanol (Sigma-Aldrich) and appropriate differentiation cytokines and antibodies for 3 to 4 days. For Th17 differentiation, differentiation media included rhTGF-ß (0.5 ng/mL), rmIL-23 (10 ng/mL), rmIL-6 (40 ng/mL), anti-IL4 (10 µg/mL), and anti-IFNγ (10 µg/mL). For Th2 differentiation, differentiation media included rmIL-4 (10 ng/mL) and anti-IFNγ (10 µg/mL), and cells were rested in rh-IL-2 (50 IU/mL) for 1-2 days prior to assay.

### Seahorse extracellular flux assays

Seahorse extracellular flux assays were performed on the Seahorse XFe96 Analyzer (Agilent) using the Seahorse XFe96 Extracellular Flux Assay Kits (Agilent). Cell culture plates were coated with Cell-Tak solution (22.4 µg/mL, Corning) overnight at 4°C before seeding the cells. Cells were plated at 150,000 cells per well with 4-5 technical replicates per sample. The GlycoStress Test (Agilent) was performed per manufacturer’s instructions. The MitoStress Test (Agilent) and Substrate Oxidation Assay (Agilent) was performed per manufacturer’s instructions except oligomycin A was used at 1.5 µM, FCCP at 1.5 µM, and rotenone/antimycin A at 0.5 µM concentrations. For the Substrate Oxidation Assay, CB-839 (MedChemExpress, 10 µM) was used to determine the acute effects of glutaminase inhibition on mitochondrial respiration. All assays were normalized using cell counts acquired via bright field imaging using a Cytation 5 Imager (BioTek). All endpoints were calculated per manufacturer’s instructions or as explained in supplemental figure 4 and in the text.

### Transmission electron microscopy (TEM)

Th17 cells were differentiated as described above from male and *Ar^Tfm^* male mice. On day 3, cells were washed with warm PBS two times and fixed in 2.5**%** glutaraldehyde in 0.1 M cacodylate for 1 hour at room temperature followed by 24 hours at 4°C. Cells were postfixed in 1% OsO4 and *en bloc* stained with 1% uranyl acetate followed by dehydration in a graded ethanol series. Samples were then gradually permeated with Quetol 651 based Spurr’s resin with propylene oxide as a transition solvent. The resin was polymerized at 60°C for 48 hours and blocks were sectioned using a Leica UC7 ultramicrotome at 70 nm nominal thickness.

Samples were stained with 2% uranyl acetate and lead citrate. TEM was performed using a Tecnai T12 operating at 100 kV with an AMT NanoSprint CMOS camera using AMT imaging software for single images.

### Mass spectrometry targeted metabolomics

Five million differentiated Th17 cells were pelleted and lysed by 5 freeze-thaw cycles. Polar metabolites were extracted with 500 μL methanol with the internal standards (13C2-Tyr and 13C3-Lactate, Cambridge Isotope Lab, Andover MA). Supernatants were evaporated to dryness under a gentle stream of nitrogen gas, and reconstituted in 100 μL of acetonitrile/water (2:1).

Metabolites were then measured by a targeted HILIC-MS/MS method developed at the Vanderbilt Mass Spectrometry Research Center (108 individual metabolites). Individual reference standards of all analytes were infused into the mass spectrometer for the optimization of ESI and selected reaction monitoring (SRM) parameters. LC-MS analysis was performed using an Acquity UPLC system (Waters, Milford, MA) interfaced with a TSQ Vantage quadrupole mass spectrometer (Thermo Scientific, San Jose, CA). Pathway analysis and statistical significance was measured using MetaboAnalyst 5.0 (Pang *et al*., 2021).

### CRISPR Screening

CRISPR screen library design and preparation were completed as previously described (Shalem et al., 2014; Sugiura *et al*., 2022; Toffalini et al., 2009). Sequences for gRNA library were chosen from the Brie library and four gRNA sequences per gene were selected along with ten non-targeting controls (NTCs), flanked by adaptor sequences, and purchased as an oligo pool from Twist Bioscience. The glutamine library contained 32 total gene targets with 138 gRNAs. Using Gibson Assembly Master Mix, oligos were cloned with the pMx-U6-gRNA-BFP vector. Plat-E retroviral packaging cells were transfected with gRNAs to package DNA into retrovirus. Naïve T cells were isolated from male and female OTII-Cas9 mice and differentiated as above except for T cell activation occurring through presentation of OVA peptide 323-339 (Sigma-Aldrich) by irradiated splenocytes from wild-type mice rather than plate-bound antibodies. Two days after activation, T cells were retrovirally transduced with retronectin-coated plates (Takara Bioscience). For the *in vitro* screen, cells were collected at day 1 and day 7 post transduction and media were changed and supplemented as necessary. For the lung inflammation screen, *Rag1*^-/-^ mice were challenged as noted above with OVA protein. Lungs were harvested, digested, and processed for gRNA frequencies as above and previously described (Sugiura *et al*., 2022). All samples maintained at least 1000-fold representation of the library through all processing. FASTQ files were analyzed using the Model-based Analysis of Genome-wide CRISPR/Cas9 Knockout (MAGeCK v0.5.0.3) method to determine statistically significant gRNA enrichments or depletions (Li et al., 2014b).

### Real-time PCR assay

Cells were cultured as stated above and washed with PBS. Total RNA was extracted using the RNeasy Mini Kit (Qiagen). cDNA was prepared using the SuperScript IV First-Strand Synthesis System (Thermo Fisher Scientific) and normalized to 400 ng of total RNA. Gene expression was determined using TaqMan primers and the TaqMan Universal Master Mix purchased from Applied Biosystems, and relative expression was normalized to *Tbp* (encoding TATA-box binding protein).

### ^18^F-Glutamine PET-CT Imaging and Glutamine Uptake Assay

Male and *Ar^Tfm^* male mice were challenged with HDM as shown in Figure 2A and described above. PET imaging studies and glutamine uptake in the cell types were performed as previously described (Reinfeld *et al*., 2021). Briefly, two hours after the last challenge, mice received a retro-orbitally injected with 1 mCi of ^18^F-glutamine synthesized at Vanderbilt University Medical Center (Hassanein et al., 2016) and rested for 40 minutes in plate-warmed cages. Mice were anesthetized using 2% isofluorane and imaged using an Inveon microPET (Siemens Preclinical) for 20 minutes. Raw data was binned as previously described and images were reconstructed into transaxial slices using the MAP algorithm. Immediately after the PET scan, the mice underwent a computed tomography (CT) scan in a NanoSPECT/CT (Mediso) and the CT images were reconstructed as previously described. Twenty-four hours after their last challenge, the mice were again injected with 1mCi of ^18^F-glutamine and allowed to rest for 40 minutes. Mice were euthanized and lungs and spleens were collected, digested, and processed as described above to obtain a single cell suspension. Cells were resuspended in MACS buffer (PBS with 2% FBS and 2 mM EDTA) and counted using trypan blue with the TC20 Automated Cell Counter (BioRad). Cell suspensions were fractionated using magnetic selection with the following: CD11b positive selection (Miltenyi Biotec) using LD columns (Miltenyi Biotec) for depletion followed by CD4 positive selection using LS columns (Miltenyi Biotec) for isolation on a Miltenyi QuadroMACS Separator according to manufacturer’s instructions. Fractions were resuspended in 1 mL medium and used for cell counts using trypan blue, flow cytometry for fraction composition, and radioactivity measurement using the Hidex Automatic Gamma Counter. To determine per cell ^18^F-glutamine activity, time-normalized counts per minute (CPM) measured on the Gamma Counter were divided by the number of viable cells as determined by the trypan count.

### SCENITH

PBMCs were thawed quickly as described in the CYTOF section for lymph nodes. Cells were then plated at 1 million cells per well in a 24-well non-tissue culture treated plate in IMDM with GlutaMAX and HEPES, 10% FBS, 1mM Sodium Pyruvate, 0.1nM Non-Essential Amino Acids, and 1% penicillin/streptomycin and activated with the Human T Cell Activation/Expansion Kit (Miltenyi) containing antibodies to CD3, CD28, and CD2 overnight at 37°C with 5% CO_2_. Bead to cell ratio was adjusted to 1:20. Cells were then plated in 96-well non-tissue culture treated U bottom plates at 400,000 cells per well in 100 µL media and allowed to rest for 1 hour at 37°C with 5% CO_2_. Cells were then treated with appropriate inhibitors or vehicle for 30 minutes (DMSO, Sigma-Aldrich) as follows: 2-deoxyglucose (100 mM, Cayman Chemical), oligomycin (1.5 µM, Cayman Chemical), and/or V9302 (10 µM, H. Charles Manning) (Schulte *et al*., 2018). After 30 minutes, cells were treated with puromycin (10 µM, Santa Cruz Biotechnology) for 40 minutes. Cells were then washed with PBS and stained with Live Dead Aqua fixable viability dye for 20 minutes at 4°C. Subsequently, cells were washed with PBS with 3% FBS and stained for surface markers 4°C as indicated. Cells were washed again with PBS with 3% FBS and fixed using the Foxp3 Transcription Factor Fix/Perm Kit (Thermo Fisher Scientific) per manufacturer instructions. Cells were washed with Perm Buffer and stained with an anti-puromycin antibody (Miltenyi Biotec) at 1:1000 at 4°C. Finally, cells were washed with Perm Buffer and PBS with 3% FBS and flow cytometry was run using a MACSQuant 16 (Miltenyi Biotec). Analysis was done using FlowJo and as previously described (Argüello *et al*., 2020).

### Statistics

All statistical analysis was performed using GraphPad Prism 9 unless otherwise indicated. Data are represented as mean ± SEM. Outliers were identified using the GraphPad Prism 9 ROUT method with a Q of 0.5% (alpha of 0.005). Comparisons between two groups were made using a two-tailed Student’s t test or Mann-Whitney U test for parametric or non-parametric data, respectively. Data were significant when *p* was less than 0.05. For targeted metabolomics, pathway analysis was conducted using MetaboAnalyst 5.0, utilizing pathway enrichment analysis with the GlobalTest and adjusted for multiple testing and the false discovery rate (FDR) and pathway impact calculated using pathway topology analysis with node importance measure determined using relative betweenness centrality. For airway hyperresponsiveness, an ANOVA of repeated measures with Bonferroni post hoc analysis was conducted. For CRISPR, statistical analysis was completed using MAGeCK (v0.5.0.3) as previously described (Li *et al*., 2014b).

Files were median normalized by read count and p values were generated using a negative binomial model comparing gRNA abundance across samples (gRNA frequency of CD4+ T cells in the lung on day 8 post-transfer or gRNA frequency on day 7 post-transduction in culture compared to gRNA frequency of CD4+ cells on day 1 post-transduction).

## Supplemental Item Titles and Legends

Table S1. Patient demographics for CYTOF on lymph nodes. Related to Figure 1 and Figure S1.

Table S2. CYTOF panel. Related to Figure 1 and Figure S1.

Table S3. Metabolic pathway analysis on Th17 cells from ArTfm male mice compared to wild-type male mice. Related to Figure 4.

Table S4. Patient demographics for SCENITH on PBMCs. Related to Figure 7 and Figure S7.

## Supplemental Figure Titles and Legends

**Figure S1.**
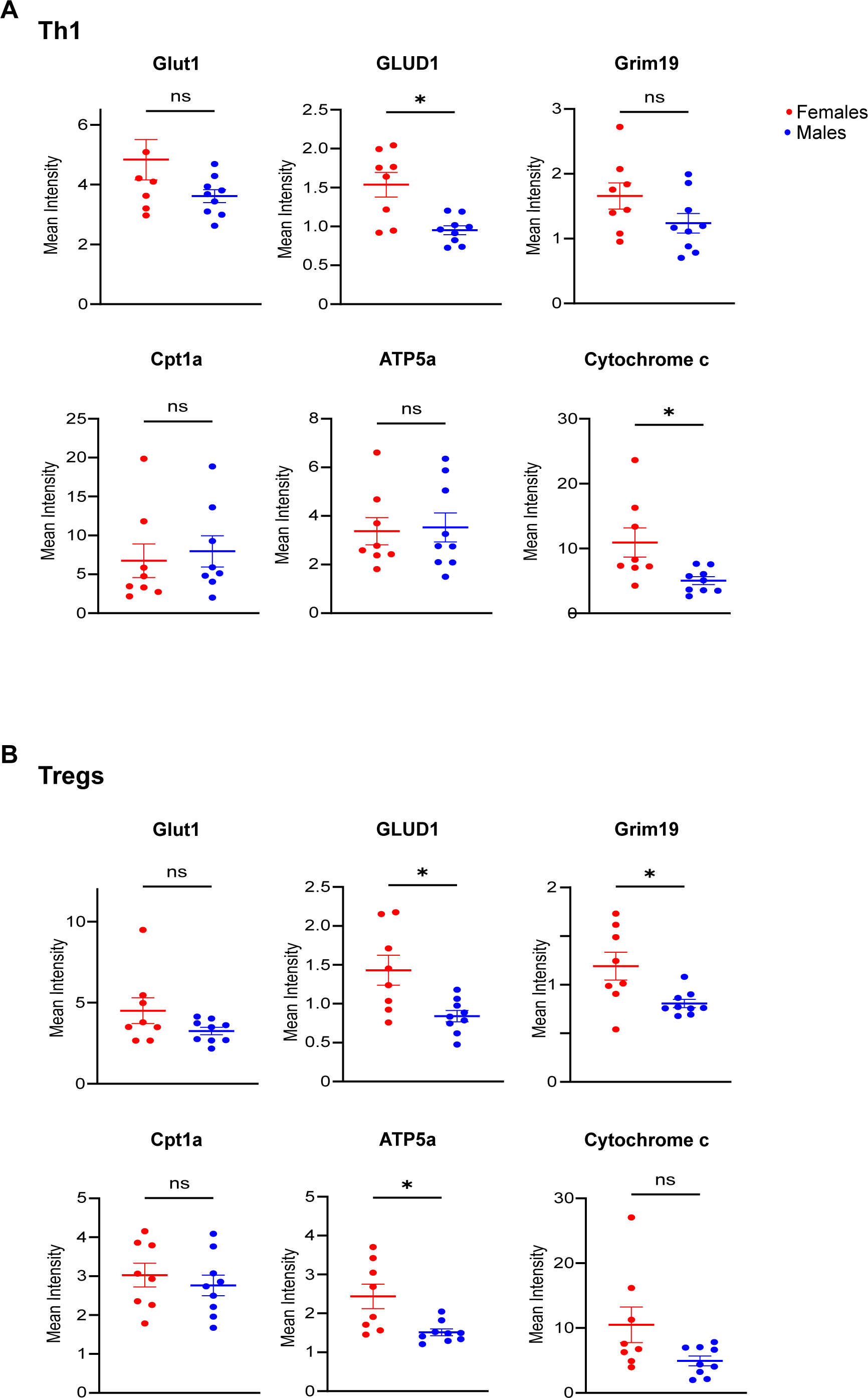
Sex differences in metabolic marker expression in Th1 and Treg. Related to. Figure 1. CYTOF was conducted on human lung draining lymph nodes from female (n=8) and male (n=9) deceased donors. **A.** Quantification of mean expression of metabolic markers on Th1 cells (gated on Live, CD3+, CD4+, CXCR5-, CXCR3+, CD25-, CCR4-). **B.** Quantification of mean expression of metabolic markers on Treg cells (gated on Live, CD3+CD4+, CXCR5-, CXCR3-, CD25+, CCR4-). All graphs show mean ± SEM. *p<0.05, ns: not significant, two-tailed Mann-Whitney U test.

**Figure S2.**
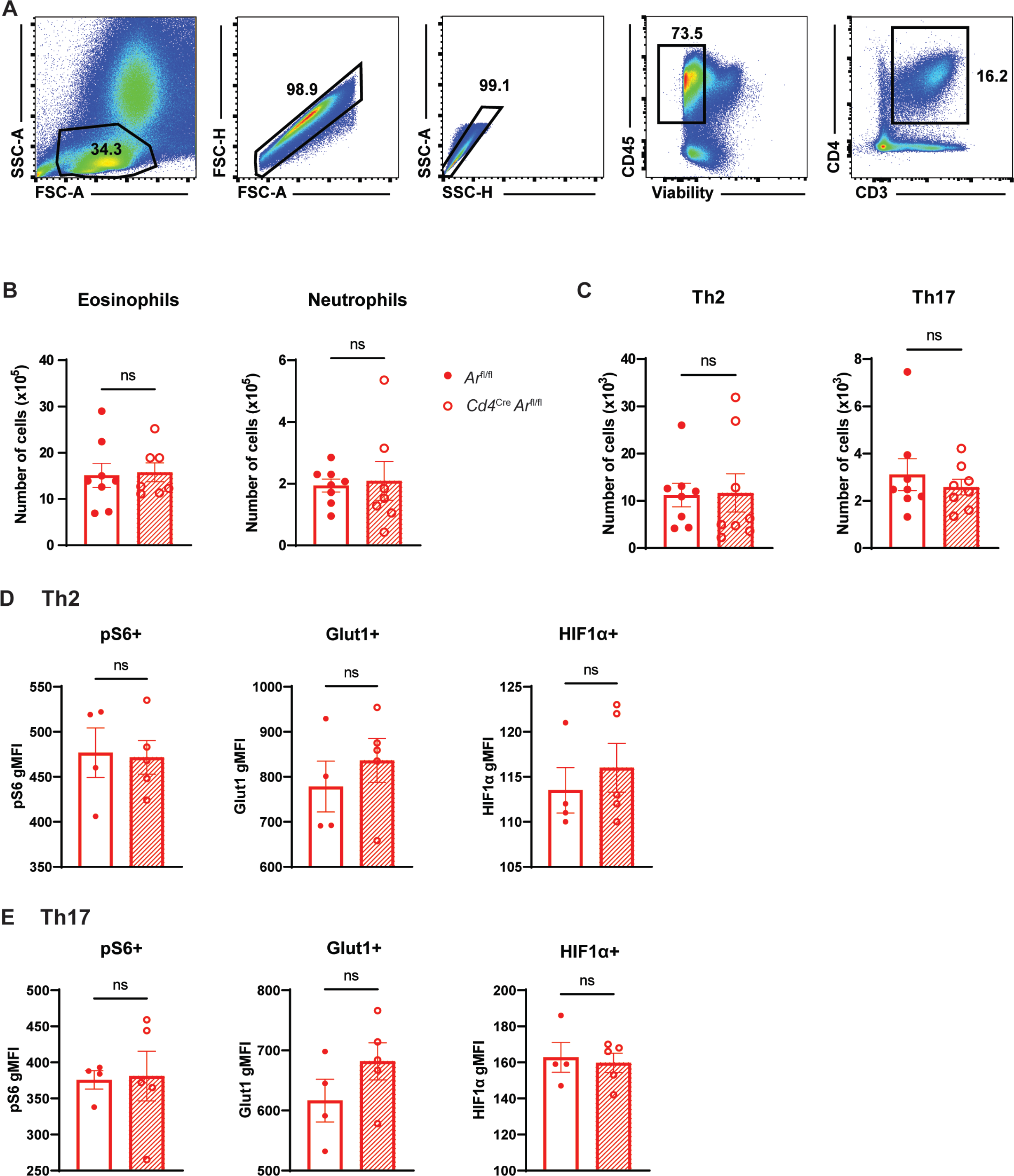
AR signaling on CD4+ cells in females does not impact airway inflammation. Related to. Figure 2**. A**. Flow gating for lung CD4+ T cells. **B**. Quantification of eosinophils and neutrophils in the BAL fluid from mice after HDM challenge (n=8 mice per group, BAL could not be collected on one mouse). **C.** Quantification of Th2 and Th17 cells in the lung after HDM challenge (n=8 mice per group). **D-E.** Expression of pS6, HIF1α **(D)** and Th17 **(E)** cells after HDM challenge (n=4-5 mice per group, representative of 2 experiments). All graphs show mean ± SEM. ns: not significant, two-tailed unpaired t test.

**Figure S3.**
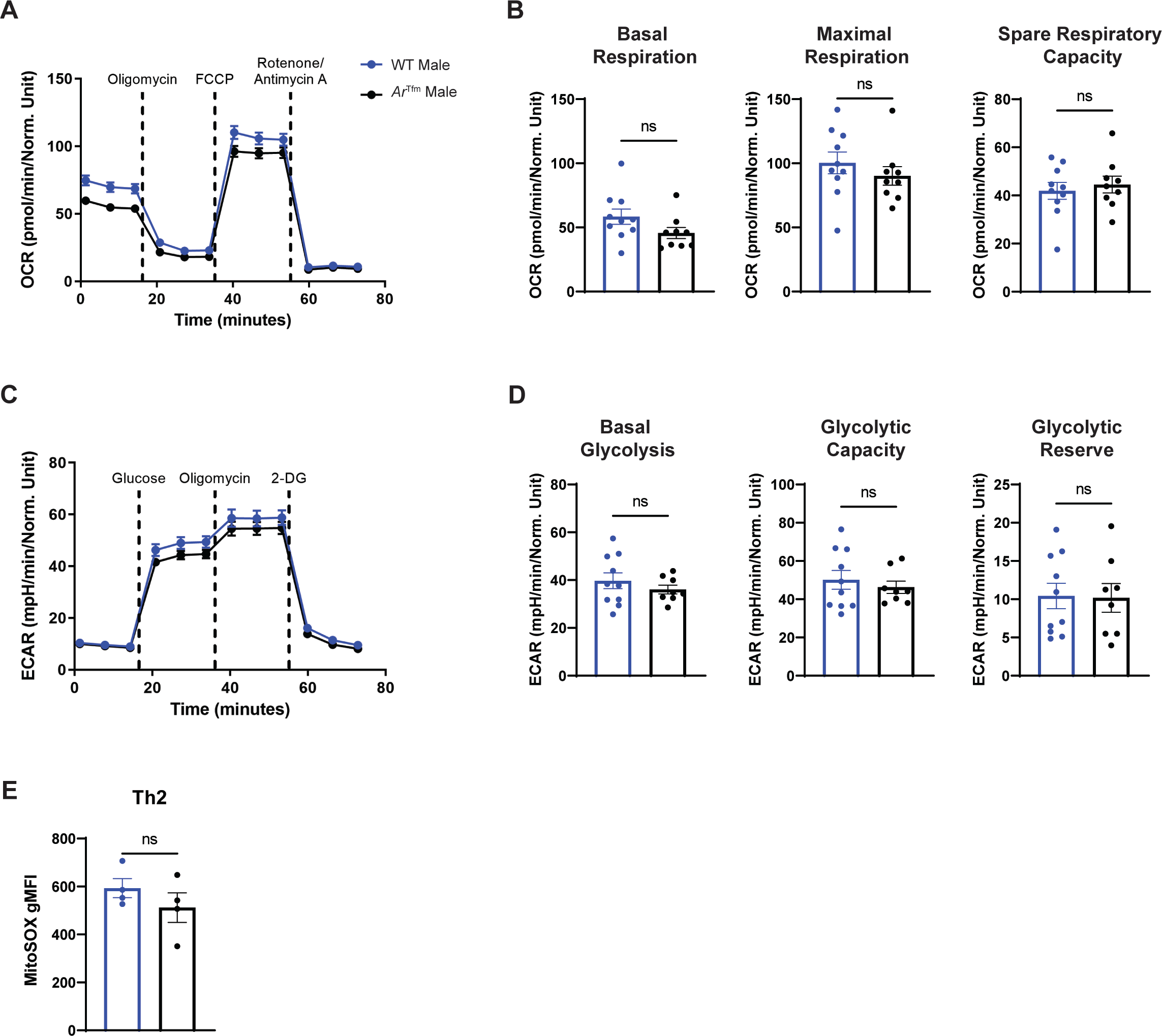
AR signaling does not impact Th2 metabolism. Related to. Figure 3**. A.** Seahorse MitoStress Test on differentiated Th2 cells from WT male and *Ar^Tfm^* male mice measures mitochondrial respiration using oxygen consumption rate (OCR) (n=9-10 mice per group). **B**. Quantified measures of basal respiration, maximal respiration, and spare respiratory capacity from panel A. **C.** Seahorse GlycoStress test on differentiated Th2 cells from wild-type male and *Ar^Tfm^* male mice to measure glycolysis using extracellular acidification rate (ECAR) (n=8 mice per group). **D**. Quantified measures of basal glycolysis, glycolytic capacity, and glycolytic reserve from panel C. **E.** Expression of MitoSOX Red, a mitochondrial superoxide marker, in differentiated Th2 cells from WT male and *Ar^Tfm^* male mice (n=4 mice per group, representative of 2 independent experiments). All graphs show mean ± SEM, ns not significant, two-tailed unpaired T test.

**Figure S4.**
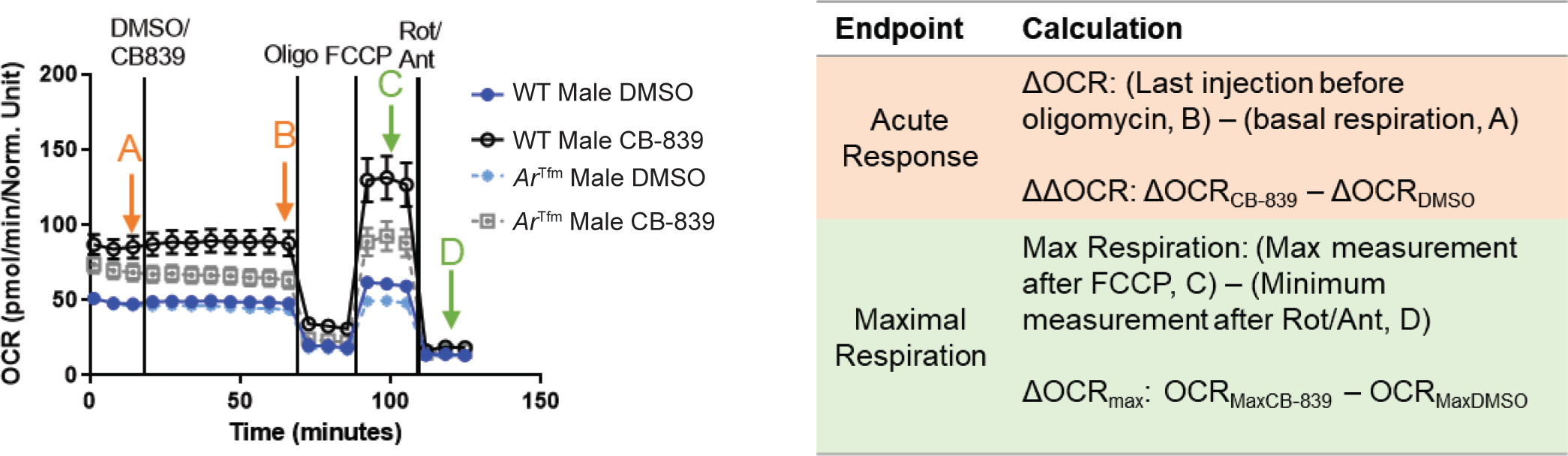
Calculations for Substrate Oxidation Assay. Related to Figure 4.

**Figure S5.**
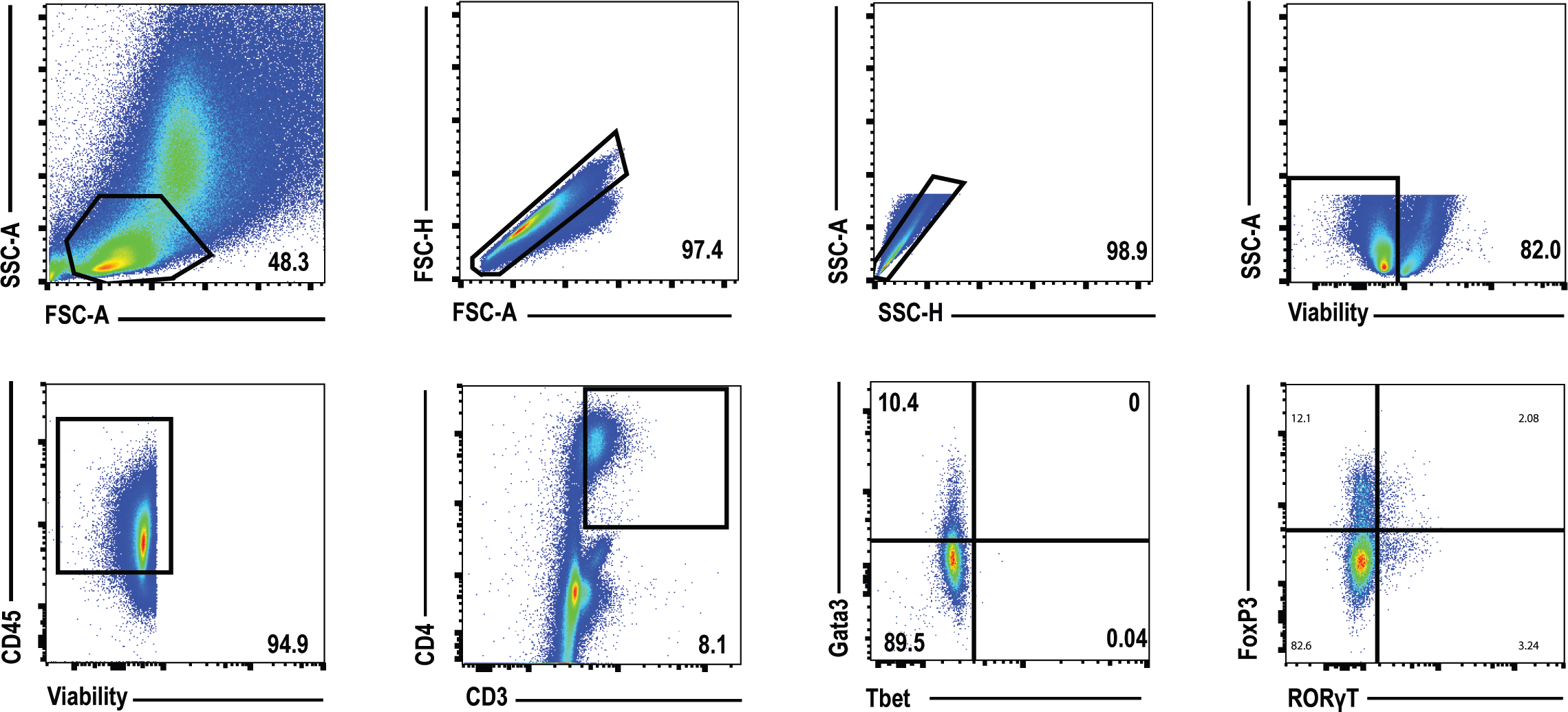
Flow gating for Th2 and Th17 cells in HDM-challenged male and female *Gls^fl/fl^* and *Cd4^Cre^ Gls^fl/fl^* mice. Related to Figure 5.

**Figure S6.**
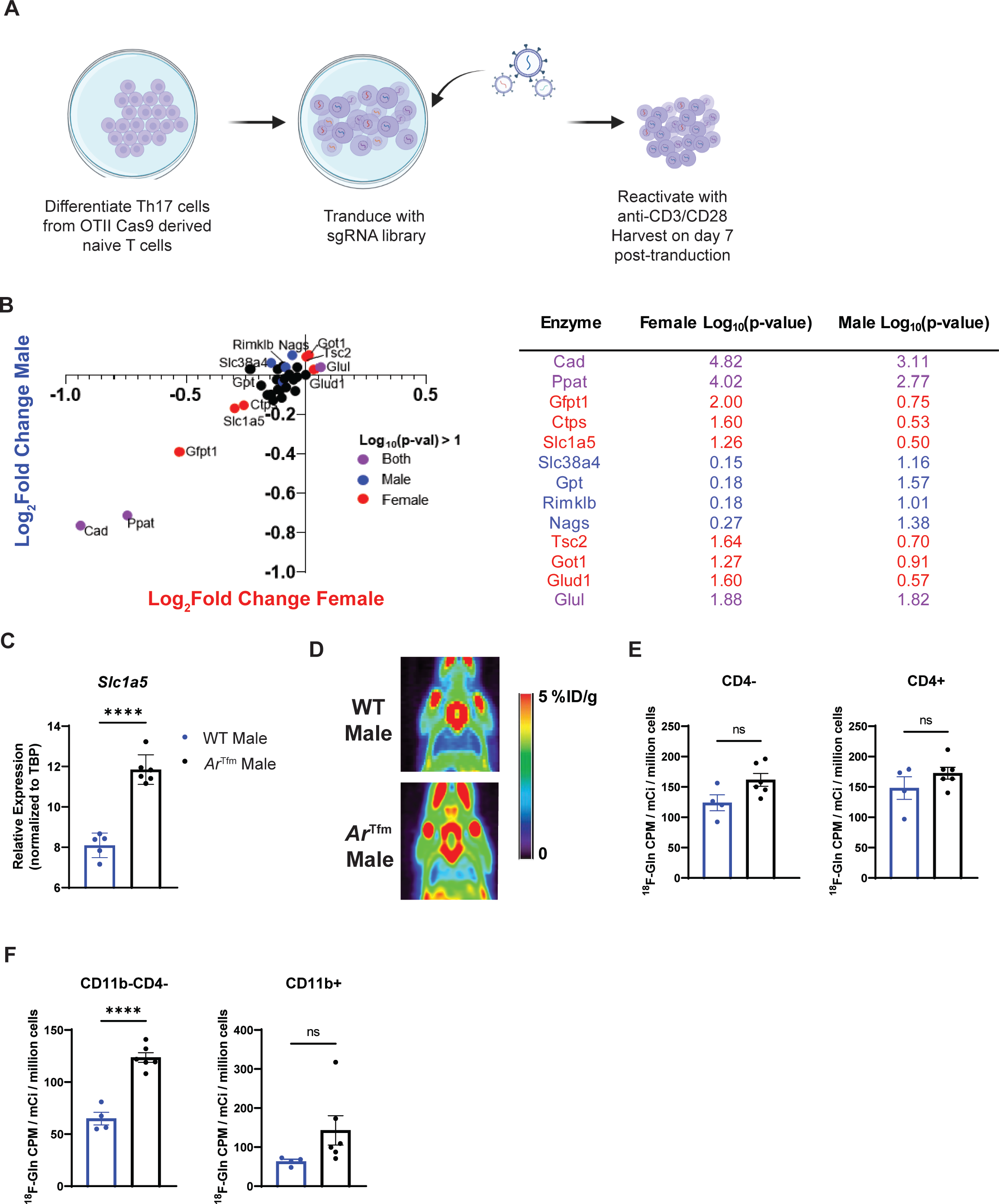
AR signaling reduces glutamine uptake in T cells. Related to. Figure 6**. A.** Model of targeted CRISPR screen using a glutamine library on differentiated Th17 cells from male and female OT-II Cas9 mice in an *in vitro* culture model. **B.** Change in gRNA abundance in lung Th17 cells from glutamine metabolism targeted CRISPR screen after OVA-induced lung inflammation with table showing statistics (n=4-5 mice, statistical analysis by MAGeCK, significantly affected represented by color of dot as shown in legend). **C.** Relative expression of *Slc1a5* in differentiated Th17 cells from wild-type male and *Ar^Tfm^* male mice (n=5-6 mice per group). **D.** Images of ^18^F-glutamine by PET Imaging. **E.** Quantification of ^18^F-Glutamine concentration in spleen CD4- and CD4+ cells by magnetic separation and gamma counting, normalized to viable cells (n=4-6 mice per group). **F.** Quantification of ^18^F-Glutamine concentration in lung CD11b-CD4- cells and CD11b+ cells by magnetic separation and gamma counting, normalized to viable cells (n=4-6 mice per group). All graphs show mean ± SEM. ****p<0.0001, ns: not significant, two-tailed unpaired T test unless otherwise specified.

**Figure S7.**
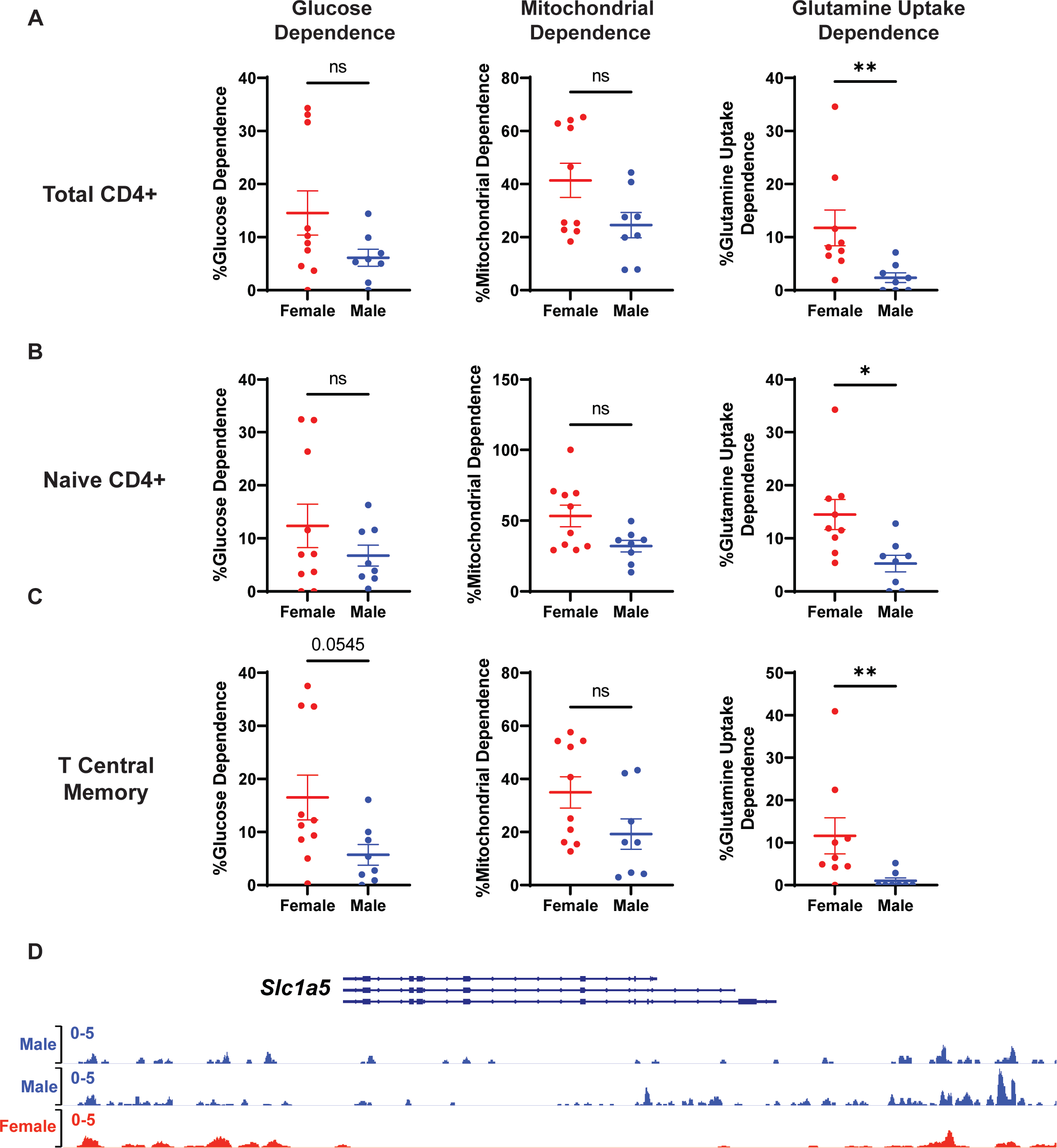
Males with severe asthma have decreased dependence upon glutamine uptake in circulating CD4+ T cell subsets compared to females with severe asthma. Related to Figure 7. A-C. Glucose dependence, mitochondrial dependence, and glutamine uptake dependence in males and females measured using SCENITH inhibitors (2-deoxyglucose, oligomycin, and V9302, respectively) in total CD4+ T (defined as Live, CD3+, CD4+, CD8-). (B), Naive CD4+ T effector memory cells (defined as Live, CD3+, CD4+ CD8-, CD45RA+, CCR7+), and (C) T central cells (defined as Live, CD3+, CD4+, CD8-, CD45RA-, CCR7+). D. H3K27Me3 MINT CHIP Seq on circulating T cells from 2 adult males and 1 adult female from the ENCODE database. All graphs show mean ± SEM. *p<0.05, **p<0.01, ns: not significant, or p-value as shown; two-tailed Mann-Whitney U test.

