## Supplemental Tables for "Androgen Signaling Restricts Glutaminolysis to Drive Sex-Specific Th17 Metabolism"

**Table S1. Patient demographics for CYTOF on lymph nodes. Related to Figure 1 and Figure S1.**

| Characteristic | Females | Males |
|---|---|---|
| Number | 8 | 9 |
| Age (y) | 30.9 ± 9.0 | 36.1 ± 9.4 |
| Race | | |
| White | 6 | 5 |
| Black/African-American | 2 | 3 |
| Asian | 0 | 1 |
| Ethnicity | | |
| Non-Hispanic | 4 | 2 |
| Hispanic | 0 | 0 |
| Not Specified | 4 | 7 |

**Table S2. CYTOF panel. Related to Figure 1 and Figure S1.**

| Marker | Clone | Mass | Marker | Clone | Mass |
|---|---|---|---|---|---|
| CD45 | HI30 | 89 | CXCR3 | G025H7 | 156 |
| Dead Rhodium | - | 103 | CD137 | 4B4-1 | 158 |
| CD66b | 80H3 | 108 | CCR7 | G043H7 | 159 |
| CD8a | RPA-T8 | 109 | CTLA4 | 14D3 | 161 |
| CD16 | 3G8 | 110 | Glut1 | NB110-39113 | 163 |
| CD14 | M5E2 | 111 | CD95 | DX2 | 164 |
| CD4 | RPA-T4 | 112 | CD44 | BJ18 | 166 |
| CD19 | HIB19 | 113 | CD38 | HIT2 | 167 |
| CD3 | UCHT1 | 116 | CYTOC | 6H2.B4 | 168 |
| CD45RO | UCHL1 | 141 | CD25 | 2A3 | 169 |
| CPT1a | 8F6AE9 | 142 | CD45RA | HI100 | 170 |
| CD127 | A019D5 | 143 | CXCR5 | RF8B2 | 171 |
| ATP5a | 7H10BD4F9 | 144 | CD57 | HCD57 | 172 |
| GRIM19 | 6E1BH7 | 145 | CXCR4 | 12G5 | 173 |
| CD20 | 2H7 | 147 | HLA-DR | L243 | 174 |
| CD27 | L128 | 148 | PD-1 | EH12.2H7 | 175 |
| CCR4 | 205410 | 149 | CD56 | CMSSB | 176 |
| CD134 | ACT35 | 150 | Cells Iridium | - | 191 |
| ICOS | C398.4A | 151 | Cells Iridium | - | 193 |
| TCRγδ | 11F2 | 152 | CD11b | ICRF44 | 209 |
| GLUD1 | polyclonal | 154 | | | |

**Table S3. Metabolic pathway analysis on Th17 cells from $Ar^{Tfm}$ male mice compared to wild-type male mice. Related to Figure 4.**

| Pathway | FDR-adjusted p-value | Impact |
|---|---|---|
| Aminoacyl-tRNA biosynthesis | 0.0131 | 0 |
| Arginine and proline metabolism | 0.0131 | 0.42 |
| Glycerophospholipid metabolism | 0.0131 | 0.03 |
| Histidine metabolism | 0.0131 | 0.22 |
| Taurine and hypotaurine metabolism | 0.0131 | 0.43 |
| Primary bile acid synthesis | 0.0131 | 0.02 |
| Arginine biosynthesis | 0.0131 | 0.25 |
| Alanine, aspartate, and glutamate metabolism | 0.0131 | 0.62 |
| D-glutamine and D-glutamate metabolism | 0.0131 | 0.50 |
| Glyoxylate and dicarboxylate metabolism | 0.0131 | 0.00 |
| Nitrogen metabolism | 0.0131 | 0.00 |
| Beta-Alanine metabolism | 0.0164 | 0.00 |
| Pantothenate and CoA biosynthesis | 0.0256 | 0.01 |
| Glycine, serine, and threonine metabolism | 0.0256 | 0.05 |
| Porphyrin and chlorophyll metabolism | 0.0262 | 0.00 |
| Phenylalanine, tyrosine, and tryptophan biosynthesis | 0.0265 | 1.00 |
| Phenylalanine metabolism | 0.0265 | 0.36 |
| Lysine degradation | 0.0268 | 0.00 |
| Valine, leucine, and isoleucine degradation | 0.0294 | 0.00 |
| Valine, leucine, and isoleucine biosynthesis | 0.0294 | 0.00 |

**Table S4. Patient demographics for SCENITH on PBMCs. Related to Figure 7 and Figure S7.**

| Characteristic | Females | Males |
|---|---|---|
| Number | 10 | 8 |
| Age (y) | 36.5 ± 5 | 33.1 ± 9.9 |
| Race | | |
| White | 2 | 5 |
| Black/African-American | 4 | 3 |
| Asian/Pacific-Islander | 2 | 0 |
| Unknown | 2 | 1 |
